# Mutualism at the leading edge: Insights into the eco-evolutionary dynamics of host-symbiont communities during range expansion

**DOI:** 10.1101/2023.04.21.537788

**Authors:** Maria M Martignoni, Rebecca C Tyson, Oren Kolodny, Jimmy Garnier

**Affiliations:** Department of Ecology, Evolution and Behavior, A. Silberman Institute of Life Sciences, Faculty of Sciences, Hebrew University of Jerusalem, Israel; Department of Mathematics, University of British Columbia, British Columbia, Canada; Laboratory of Mathematics, CNRS, Université Savoie-Mont Blanc, Université Grenoble Alpes, Chambery, France

**Keywords:** eco-evolutionary dynamics, mathematical model, mutualism, range expansion, arbuscular mycorrhizal fungi, host-microbial communities, differential equations

## Abstract

The evolution of mutualism between host and symbiont communities plays an essential role in maintaining ecosystem function and should therefore have a profound effect on their range expansion dynamics. In particular, the presence of mutualistic symbionts at the leading edge of a host-symbiont community should enhance its propagation in space. We develop a theoretical framework that captures the eco-evolutionary dynamics of host-symbiont communities, to investigate how the evolution of resource exchange may shape community structure during range expansion. We consider a community with symbionts that are mutualistic or parasitic to various degrees, where parasitic symbionts receive the same amount of resource from the host as mutualistic symbionts, but at lower cost. The selective advantage of parasitic symbionts over mutualistic ones is strengthened with resource availability (i.e. with host density), promoting mutualism at the range edges, where host density is low, and parasitism in the population core, where host density is higher. This spatial selection also influences the speed of spread. We find that the host growth rate (which depends on the average benefit provided by the symbionts) is maximal at the range edges, where symbionts are more mutualistic, and that host-symbiont communities with high symbiont density at their core (e.g. resulting from more mutualistic hosts) spread faster into new territories. These results indicate that the expansion of host-symbiont communities is pulled by the hosts but pushed by the symbionts, in a unique push-pull dynamic where both the host and symbionts are active and tightly-linked players.

## Introduction

Climate change and anthropogenic disturbance are inducing species to engage in range expansion toward more suitable habitats at an unprecedented rate. Understanding the mechanisms governing population spread is therefore a key priority in conservation biology. Nonetheless, predicting the specific contours of a species’ range expansion remains challenging (Fournier et al, 2019). Spatial spread is determined by processes acting at the edge of a species’ range, where population densities are lower in comparison to those in the core of the range (Chuang and Peterson, 2016). Low-density populations experience unique interaction structures and selective pressures, such as decreased intraspecific competition, or altered reproductive success (Phillips, 2009; Huang and Peng, 2016)), that together favour the evolution of certain traits with respect to others and can affect population spread in an unexpected manner.

The range expansion of a host population associated with a mutualistic symbiont community represents a particularly challenging study system. The term ‘host-associated community’ refers to a community of small organisms with a short generation time (the ‘symbionts’) associated with a large host (Bronstein, 2015). Host-associated communities may encompass a collection of parasitic, commensal, and/or mutualistic organisms where, to be defined as ‘mutualistic’ the associated symbionts should have an overall positive effect on host fitness, at least to some extent (Bronstein, 2015). If the fitness of a host species is affected by the presence of its mutualistic symbionts, it follows that the spatial spread of the host population will be intrinsically coupled to the density and characteristics of its associated symbionts at the range edge. Additionally, lower population densities of the host at the range edges of an expanding population may lead to the selection of symbionts with mutualistic traits that differ from those observed in the population core, further affecting the expansion dynamics (Koella, 2000; Van Dyken et al, 2013). Mutualism thus adds a further degree of complexity to the evaluation of the speed of spread of hosts and symbionts.

Host-associated mutualistic communities are widespread in nature, and many have well-established ecological and economic significance. Examples include the crucial role played by mycorrhizal fungi in plant growth and agricultural productivity (Smith and Read, 2010), or the fundamental functions performed by zooxanthellae in supporting coral reef ecosystems and fisheries (Muller-Parker et al, 2015). Only recent studies, however, have investigated how mutualistic interactions can affect the range limits of a host population and its symbionts (Afkhami et al, 2014; Araújo and Rozenfeld, 2014; Stanton-Geddes and Anderson, 2011; Dickie et al, 2017; Godsoe et al, 2017; Fournier et al, 2019; Benning and Moeller, 2021; Paquette and Hargreaves, 2021; Stephan et al, 2021). In this context, much greater attention has been given to the influence of abiotic factors (Chen et al, 2011; Spence and Tingley, 2020), or to the negative impacts of biotic factors such as predation, competition, or pathogen transmission (Phillips et al, 2010; Brown and Vellend, 2014; Wan et al, 2016). Theoretical approaches have also been scarce (Case et al, 2005; Brooker et al, 2007; Mack, 2012; Kubisch et al, 2014; Shaw, 2022), with very few mathematical models treating hosts and their multiple symbionts as separate entities capable of influencing each others’ population dynamics and evolutionary trajectories.

Even less is known about how the interaction dynamics between hosts and symbionts are shaped by spatial dispersal (Mony et al, 2022; Hu et al, 2022), and whether the community of symbionts in a core population should be expected to differ from what is observed at the edges of the expansion range (Doebeli and Knowlton, 1998). Empirical work on host-microbial symbioses has revealed changes in microbial diversity at the range edge (Lankau and Keymer, 2016; Fowler et al, 2023). This variation could lead to a decrease in host fitness or could help the host to better adapt to new environmental challenges, depending on the microbial composition observed in the edge community. These studies, however, do not address the question of whether range expansion favours selection for specific mutualistic traits in the symbionts, thereby affecting the structure of the symbiotic community. Currently, only a few studies of protective plant-ant mutualisms have tackled this issue; these studies have identified differences between the mutualistic investment of ants in the core and at the range edge (Léotard et al, 2009; Vittecoq et al, 2012).

To address these questions, we develop a theoretical framework considering the mutualistic interactions of a population of one type of plant (the ‘host’) and a community of host-associated arbuscular mycorrhizal fungi (the ‘symbionts’). We consider host and symbionts to interact by exchanging resources necessary for their growth, where access to the host’s resource is the same for all symbionts, regardless of the benefit they provide in return. We further assume that symbionts have the ability to disperse independently from the host and to transmit horizontally within the host population.

Host control, intended as any mechanism by which hosts can actively select for more beneficial symbionts in the community, has been a popular explanation for the persistence of mutualism in the face of cheaters (Hoeksema and Kummel, 2003; Bever, 2015; Bachelot and Lee, 2018; Christian and Bever, 2018). Host species have been found to have some influence over the composition and persistence of their symbionts, e.g. through gene regulation (Wier et al, 2010; Davenport et al, 2015), through secretion of compounds such as amino acids, sugars, and organic acids (Yuan et al, 2015; Frenkel and Ribbeck, 2017), or through preferential allocation of resources (Bever et al, 2009; Kiers et al, 2011). The universality of host control mechanisms, however, and the extent to which these mechanisms apply in natural contexts, have both been questioned in numerous studies, particularly in the context of the mycorrhizal symbiosis (Cameron et al, 2008; Walder et al, 2012; Walder and Van Der Heijden, 2015; Zhang et al, 2015; van der Heijden and Walder, 2016).

In the absence of host control mechanisms, vertical transmission of symbionts has also been found to favour mutualism (Ewald, 1987; Sachs et al, 2004). Indeed the faithful transmission of symbionts from parent to offspring may promote the evolution of more mutualistic traits when the reproductive success of a host and its symbionts is linked (Sachs et al, 2011; Frederickson, 2013). Symbionts, however, are often horizontally or environmentally inherited (Wilkinson and Sherratt, 2001; Vandenkoornhuyse et al, 2015; Shade et al, 2017). In this case, a symbiont may remain parasitic rather than engage in a mutualistic interaction (Ferdy and Godelle, 2005; Drew et al, 2021a), and little is known about how mutualism can persist when the fates of the symbionts and the host are decoupled. We have therefore developed a model that allows us to investigate mutualism persistence in the paired conditions of no host control and sharing of symbionts among hosts.

Evolutionary changes in symbionts have been proposed to play a key role in structuring microbial communities, and represent an important factor to consider in elucidating the interaction dynamics of a host and its symbionts (Gómez et al, 2016; Miller et al, 2018). We, therefore, assume that the mutualistic investment of each symbiont (quantified by the rate at which the symbiont provides a benefit to the host) can change in time through a diversification process, e.g., genetic or phenotypic mutation. We develop a theoretical framework in which the evolutionary dynamics of the symbionts and their ecological interactions with the host occur on the same timescale. Specifically, we ask the following questions:

i. How do eco-evolutionary dynamics shape the structure of host-associated mutualistic communities?
ii. How does symbiont investment, in the mutualistic relationship with the host, differ between the population core and the edge of the expansion range?
iii. How does resource supply of a host population toward its symbionts affect the spread of both host and symbiont populations?

Our analysis provides insights into the eco-evolutionary dynamics and linked expansion dynamics of a host population and its symbionts. To make our work more concrete, we select the mycorrhizal mutualism as our motivating example, but our model and results apply to multiple ecological systems (Palmer et al, 2003; Robinson et al, 2010).

## Models and Methods

We present an eco-evolutionary model describing the dynamics of a host population of plants supporting a host-dependent community of arbuscular mycorrhizal fungi (AMF), where the associated AMF compete amongst each other for space (Thonar et al, 2014; Engelmoer et al, 2014). The plant and AMF densities correspond to the amount of biomass (leaves, roots, hyphae, etc.) present at each time and location and are described by the variables *p*(*t, x*) and *m*(*t, x, α*), respectively. The AMF community is composed of fungi that differ in their ability *α* to deliver resources to host plants, as explained in detail below.

Individuals within the plant population and AMF community are each characterized by a trait quantifying their mutualistic investment, i.e., the rate at which each delivers resource to the partner: Each plant delivers carbon to the AMF at a rate quantified by parameter *β*, while each associated AMF delivers phosphorus to the plant at a rate quantified by parameter *α*. The mutualistic investment of the fungi is subject to evolution, i.e., *α* can evolve through mutation or recombination during spore production and spread, or during root colonization (Vandenkoornhuyse et al, 2001; den Bakker et al, 2010). Thus, a plant-dependent community of AMF includes fungi that differ in their mutualistic investment *α*. When the mutation rate is high, or mutations with large effects on traits are rare (as it is commonly observed in microbial communities (LeClerc et al, 1996; Oliver et al, 2000; Trindade et al, 2010)), diversification of the trait *α* can be modelled as a diffusion process with a diffusion or *diversification rate d*_*m*_, corresponding to the product of the mutation rate and half the variance associated with mutational effects (Fleming, 1979). In addition to trait evolution, we consider the plant population and AMF community to undergo spatial expansion through the random dispersal of seeds or spores, respectively, and we quantify their dispersal ability by the diffusion coefficients *D*_*p*_ (for the plant) and *D*_*m*_ (for the associated AMF).

We thus obtain the following model for a population of plants with biomass density *p*(*t, x*) interacting with a community of AMF with total biomass density *M* (*t, x*), composed of multiple AMF whose biomass density *m*(*t, x, α*) depends on their mutualistic investment *α*:

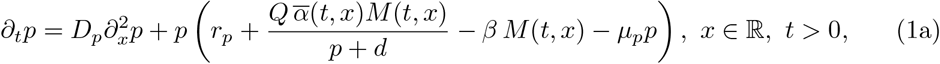

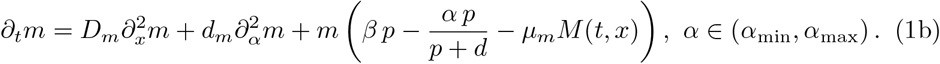

Note that *p, m*, and *M* have been converted into dimensionless quantities with arbitrary units. The parameter *α*_min_ is the minimal mutualistic investment of the AMF (which can be zero) and *α*_max_ *>* 0 is the maximal mutualistic investment. Each AMF is considered parasitic (if the cost of the symbiont for the plant is higher than the benefit it provides) and mutualistic (if the benefit provided is higher than the cost), as we will explain in detail below (see Eq. (3)). The quantity 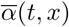 is the mean mutualistic investment of the AMF community at location *x* and time *t* and is defined as

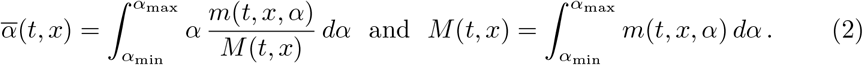

To ensure that mutations do not affect the total population biomass of the symbiont community, Eqs. (1) have no-flux boundary conditions on the boundary of the trait domain (*α*_min_, *α*_max_), that is,

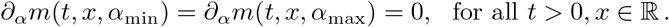

The delivery of resources from the plant to the AMF depends linearly on plant and AMF densities (see fourth and third terms in Eqs. (1a) and (1b) respectively). Resource delivery from the AMF to the plant increases linearly with increasing fungal density, and nearly linearly with increasing plant density when plant density is low (relative to parameter *d*) (see third and fourth terms in Eqs. (1a) and (1b) respectively). When plant density is large (relative to parameter *d*) resource delivery tends toward a dependency on fungal density only, as we assume host availability to not be a factor affecting the resource delivery capacity of associated symbionts (see Martignoni et al (2020) for more details about the biological assumptions underlying the choice of these functional responses for the benefit and cost of mutualism).

The nutrients (e.g., phosphorus) received by the plant from the AMF, and the carbon received by the AMF from the plant, are converted into plant and AMF biomass, respectively. The parameter *Q* represents the efficiency of these conversions (see Martignoni et al (2020)). Parameters *μ*_*p*_ and *μ*_*m*_ correspond to the density-dependent rates at which resources are directed to the maintenance of the existing plant and AMF biomass, respectively. These costs encompass, for example, energetic costs and the cost of direct competition. Note that the maintenance term for the AMF community depends on the total AMF density *M* (*x, t*) (see last term of Eq. (1b)), and therefore includes direct competition between symbionts.

Below, we summarize the main questions we investigate using our model, and the approaches we take to do so.

### The eco-evolutionary dynamics of host-associated communities

We first consider the model of Eq. (1) in the absence of spatial spread (i.e., *D*_*m*_ = *D*_*p*_ = 0). We aim to understand how a symbiotic community can emerge from natural selection and persist in association with a population of host plants. For this purpose, we investigate the coupled dynamics of a community of AMF associated with a population of host plants, and we consider the role that evolutionary changes in the mutualistic investment of the symbionts (*α*) may play in the establishment and persistence of mutualistic communities. Symbionts with low mutualistic investment reduce the growth rate of the host. We will refer to these symbionts as ‘parasitic’, and we will define them as AMF characterized by *α* ⩽ *α*_*c*_, where *α*_*c*_ is the threshold below which an obligate mutualistic plant can not survive in the presence of that single symbiont (see SI.A for mathematical details). The threshold *α*_*c*_ is defined by

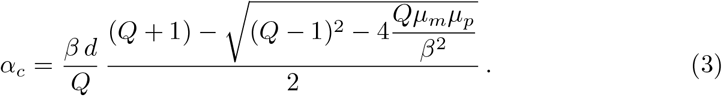

Symbionts with large mutualistic investment (*α* ⩾ *α*_*c*_) enhance the host growth rate, which allows the host-associated community to survive. We will refer to these symbionts as ‘mutualistic’. We investigate the mechanisms leading to the coexistence of mutualistic and parasitic symbionts in a community supported by a host population. In addition, we study how the mutualistic investment (*β*) of host plants toward their associated AMF affects the distribution of mutualistic traits (*α*) among the AMF community and the proportion of parasitic and mutualistic symbionts in the community.

### Community structure at the range edges

We aim to understand how dispersal structures the average mutualistic investment 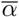 along a range expanding community of AMF associated with a population of host plants. The travelling wave solutions of Eqs. (1), i.e., solutions whose fixed profile moves at a constant speed, can be used to understand this propagation. We compare the proportion of parasitic and mutualistic symbionts and their average mutualistic investment 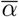 at the edge and in the core of an expanding plant-AMF community. By computing the plant and AMF growth rates along the wave front, we also gain insights into whether the expansion of host-associated communities can be defined as a pushed wave (i.e. driven by a growth rate that is highest some distance behind the wave front) or pulled wave (i.e. driven by a growth rate that is highest at the leading edge). We also investigate the ancestry of the symbionts at the leading edge, to determine the relative contribution of parasitic and mutualistic symbionts to the community at the leading edge.

### The spread of a host population and its symbionts

We aim to understand how mutualistic interactions between a plant and its associated AMF affect the speed of spread of a host and its associated symbionts in a homogeneous landscape. We first compute the speed of spread of a host-symbiont community for different values of the rate *β* at which the plant provides carbon to its AMF. We further investigate how the speed of spread is affected by the difference between the dispersal abilities of the plant and the AMF. More precisely, we vary the ratio between the dispersal ability of the plant population (*D*_*p*_) and that of the AMF community (*D*_*m*_). Finally, we test how results differ when the plant is an obligate versus a facultative mutualist (i.e., when the intrinsic growth rate of the plant *r*_*p*_ is zero, if the mutualism is obligate, or larger than zero if the mutualism is facultative).

**Table 1.** Parameters of the model with either the range of values we consider, or the default value used in the simulations.

| Parameters |  | Range |
| --- | --- | --- |
| $x$ | variable representing position in space | $(-\infty, \infty)$ |
| $\alpha$ | AMF mutualistic investment trait | $[0, 5]$ |
| $r_p$ | intrinsic growth rate of plant | $[0, 0.8]$ |
| $\beta$ | plant carbon supply rate | $[0.29, 13]$ |
| $Q$ | relative nutrient conversion efficiency | 6 |
| $d$ | half-saturation constant | 1.2 |
| $\mu_p$ | Parameters of plant density dependence | 0.3 |
| $\mu_m$ | Parameter of AMF density dependence | 0.3 |
| $d_m$ | diversification rate | $[10^{-5}, 10^{-1}]$ |
| $D_p$ | dispersal rate of plant | 0.1 |
| $D_m$ | dispersal rate of AMF | 0.1 |

## Results

### The eco-evolutionary dynamics of host-associated communities

In the absence of dispersal, our model predicts the establishment and persistence of a mutualistic host-associated community (Fig. 1(a)). The emerging community is composed of a combination of symbionts that are parasitic or mutualistic to various degree, where the mutualistic investment of the symbionts does not converge to a specific value but is distributed within the community across the range *α*_min_ to *α*_max_ (Fig. 1(b)).

**Fig. 1.**
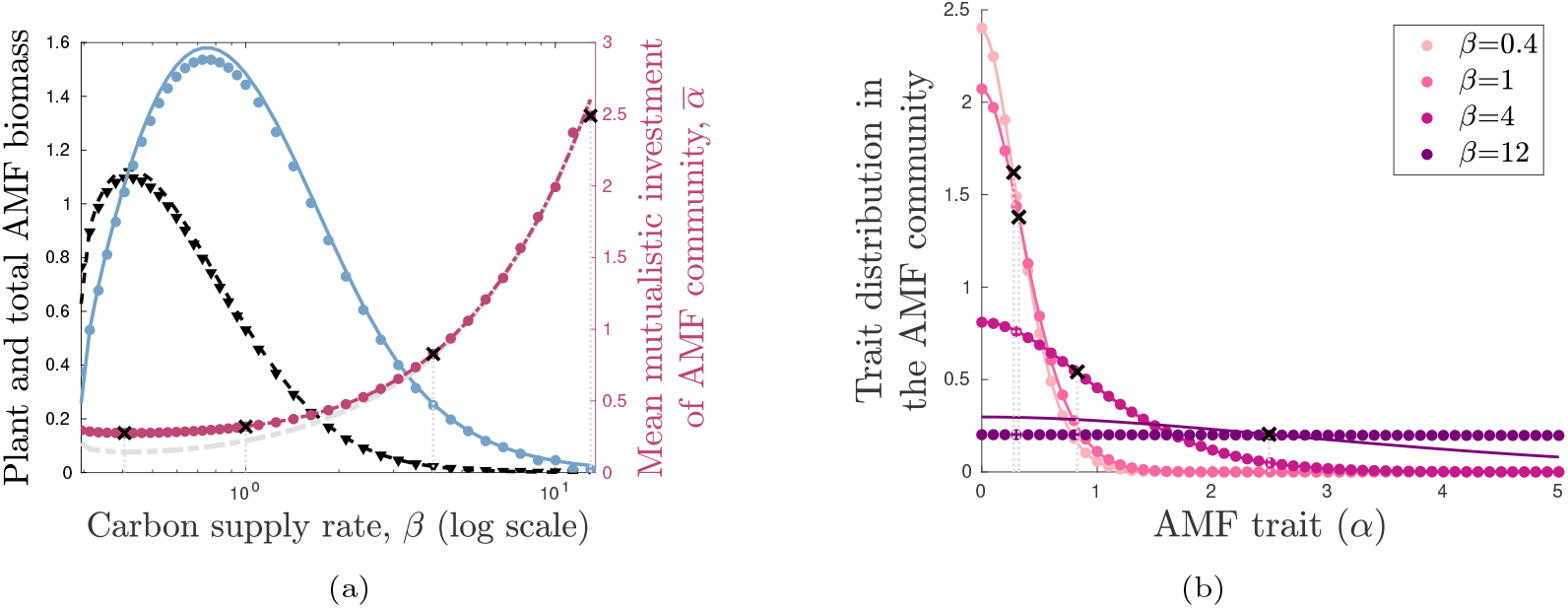
Host-symbiont community dynamics in the absence of dispersal. Panel (a), Plant density at equilibrium (black) and total density of the AMF community (blue) as functions of the plant carbon supply rate *β*. The mean mutualistic trait 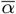 of the AMF community is shown in red and the mutualism/parasitism threshold *α*_*c*_ is shown in grey. Black asterisks correspond to the simulation results in panel (b). Panel (b), Trait distribution in the AMF community as a function of AMF phosphorus supply rate *α* and for various values of plant carbon supply rate *β*. For both panels, symbols correspond to equilibria of the model (1) without spatial spread (*D*_*p*_ = *D*_*m*_ = 0) obtained from numerical simulations, while curves correspond to theoretical approximations defined in SI.A. Parameter values used for the simulations are: *d*_*m*_ = 0.01, *Q* = 6, *μ*_*p*_ = *μ*_*m*_ = 0.3, *α*_min_ = 0 and *α*_max_ = 5.

Mutualistic symbionts receive the same benefit from the hosts as parasitic ones (proportional to *β p*) but at a higher cost (proportional to *αp/*(*p* + *d*), where *α* is lower for parasitic symbionts) as described by Eq. (1b). Parasites thus benefit from higher fitness and reach higher densities than mutualists, which confers a competitive advantage to parasitic symbionts. As a result, the trait distribution of AMF has its maximum at *α*_min_ and decreases with increasing *α* (Fig. 1(b) and SI.B for mathematical details). However, the proportion of parasites in the AMF community, that is AMF with low mutualistic investment (*α*_min_ ⩽ *α* ⩽ *α*_*c*_), is not always larger than 1*/*2. The prevalence of parasites in the AMF community truly depends on the plant carbon supply rate *β* as well as the diversification rate *d*_*m*_ (see Fig. B2).

The selection strength favouring parasitic symbionts (with lower values of *α*) increases with plant density *p* (see Eq. (1b)). However, host density decreases as symbionts become more parasitic and the mean mutualistic investment 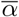 decreases (see Eq. 1a). Thus, in the absence of mutations, mutualistic symbionts are competitively excluded by parasitic symbionts, with a consequent reduction in plant growth and collapse of the whole community (see SI.A for more details). Mutations cause parasites to evolve into better mutualists, driving diversification in *α* toward higher mutualistic investment. Even when diversification occurs at an extremely low rate (Fig. B1(a) in SI.B), this process can balance selection due to competition between symbionts (which selects for low values of *α*) to produce a mean mutualistic investment of the AMF community large enough to support plant growth (Fig. 1(a)).

From the perspective of the symbionts, the intertwined effects of mutation and selection generate an average mutualistic investment 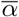 that decreases with selection strength and thus decreases with respect to plant density. More specifically, the resulting balance between selection and mutation produces a distribution of AMF density as a function of plant density *p* that can be approximated by the Airy function truncated at *α*_*max*_, with mean 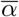 satisfying

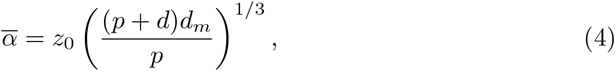

where *z*_0_ is a positive constant that depends only on the Airy function solving the dimensionless problem Ai^*′′*^(*z*) − *z* Ai(*z*) = 0 on ℝ (see SI.B.3 for more details and Fig. 1b). In addition, the mean mutualistic investment 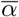 is increasing with the diversification rate *d*_*m*_ (Fig. B1).

From the perspective of the plants, the ecological interaction between the plants and the AMF community produces dynamics with antagonistic effects. Host plants benefit from the community at rate 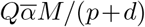, which is increasing with the average mutualistic investment 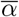, but decreasing with host density *p*. At ecological equilibrium, plant density satisfies

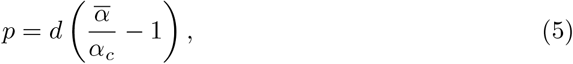

where *α*_*c*_ is the parasitic/mutualistic threshold defined by (3) (see SI. B).

The antagonistic outcomes of the evolutionary process described by (4), and the ecological interactions between the host plants and symbiont community described by (5), produce an eco-evolutionary equilibrium. The eco-evolutionary feedback between plant density and AMF mean mutualistic trait is key to the persistence of a community where parasitic and mutualistic symbionts coexist (see more details in Box 1 and SI.A).

#### Box 1

**Forces stabilizing the eco-evolutionary dynamics of host-symbiont communities**

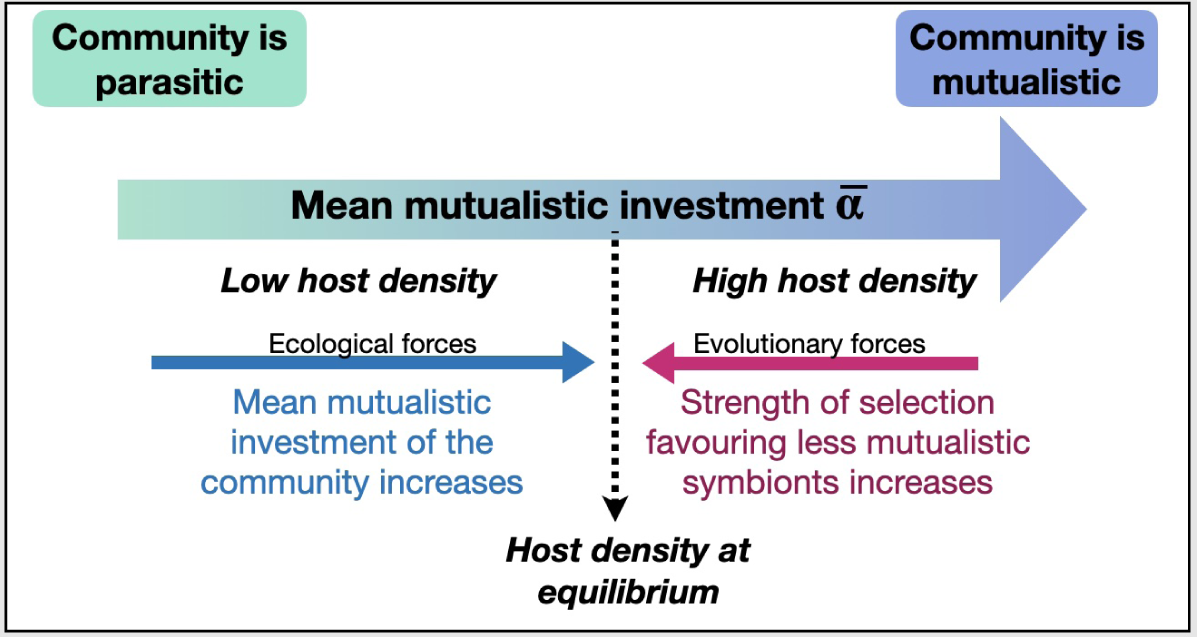

Diagram illustrating the evolutionary and ecological forces stabilizing the population dynamics of host-symbiont communities. Ecological interactions between hosts and symbionts tend to increase host density in a way that is proportional to the mean mutualistic investment 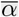 of its symbionts (see Eq.(5)). Evolutionary forces, on the other hand, tend to favor parasitic symbionts, which have a greater fitness advantage over mutualistic symbionts. This advantage increases with increasing resource availability, and thus with increasing host density (see Eq.(4)). If the community is highly parasitic, 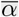 assumes a lower value, which causes a reduction in host density and a decrease in the fitness advantage of parasitic symbionts, which then leads the community to become more mutualistic on average. If the community is more mutualistic, host density increases and so does the fitness advantage of parasitic symbionts. The balance between evolutionary forces (selecting for more parasitic traits through the benefits of cheating) and ecological forces (selecting for more mutualistic traits when plant density is high) stabilizes host density around a fixed value.

Carbon supply to the symbionts (parameter *β*) represents the mutualistic investment of the host plants. It determines the growth of the associated AMF and, in turn, plant growth. Plant density at equilibrium is therefore affected by *β* (see Eq. (5)). Since the plant density affects community structure through the evolutionary dynamics of symbionts (see Eq. (4) and SI.B), the carbon supply rate *β* is also an indirect determinant of community structure. In particular, a low rate of carbon supply *β* from the host to the symbionts does not provide enough resource for the AMF community to grow and support host growth, while a large *β* leads to a high cost of mutualism for the plant population, and thus directly reduces its growth. Plant density is therefore maximized at an intermediate value of *β* (Fig. 1(a)).

### Community structure at the range edges

#### Mutualistic investment at the range edges

In this section we focus on the specific contribution of parasitic and mutualistic symbionts in shaping the expansion dynamics of the community. We observe that in the core of an expanding population, coexistence of parasitic and mutualistic symbionts is dynamically stable, with selection favouring parasitic symbionts and allowing them to grow larger than mutualistic symbionts (Fig. 2(a)), consistent with the non-spatial eco-evolutionary dynamics described above. In contrast, toward the expansion edges plant density decreases, which reduces the strength of selection acting against mutualistic symbionts and thus increases the mean mutualistic investment of the community. The resulting effect on the distribution of *α* across the AMF community is similar to the one observed when varying *β* in the absence of dispersal, but with position along the wavefront, *x*, playing the role of *β* (cfr. Fig. 1(b) and Fig. 2(b)). Indeed, the positive dependence of the selection strength on plant density, combined with the low plant density at the leading edge of the traveling wave, explains the decrease in the proportion of parasitic symbionts toward the traveling front (Fig. 2(c) and SI.B.4).

**Fig. 2.**
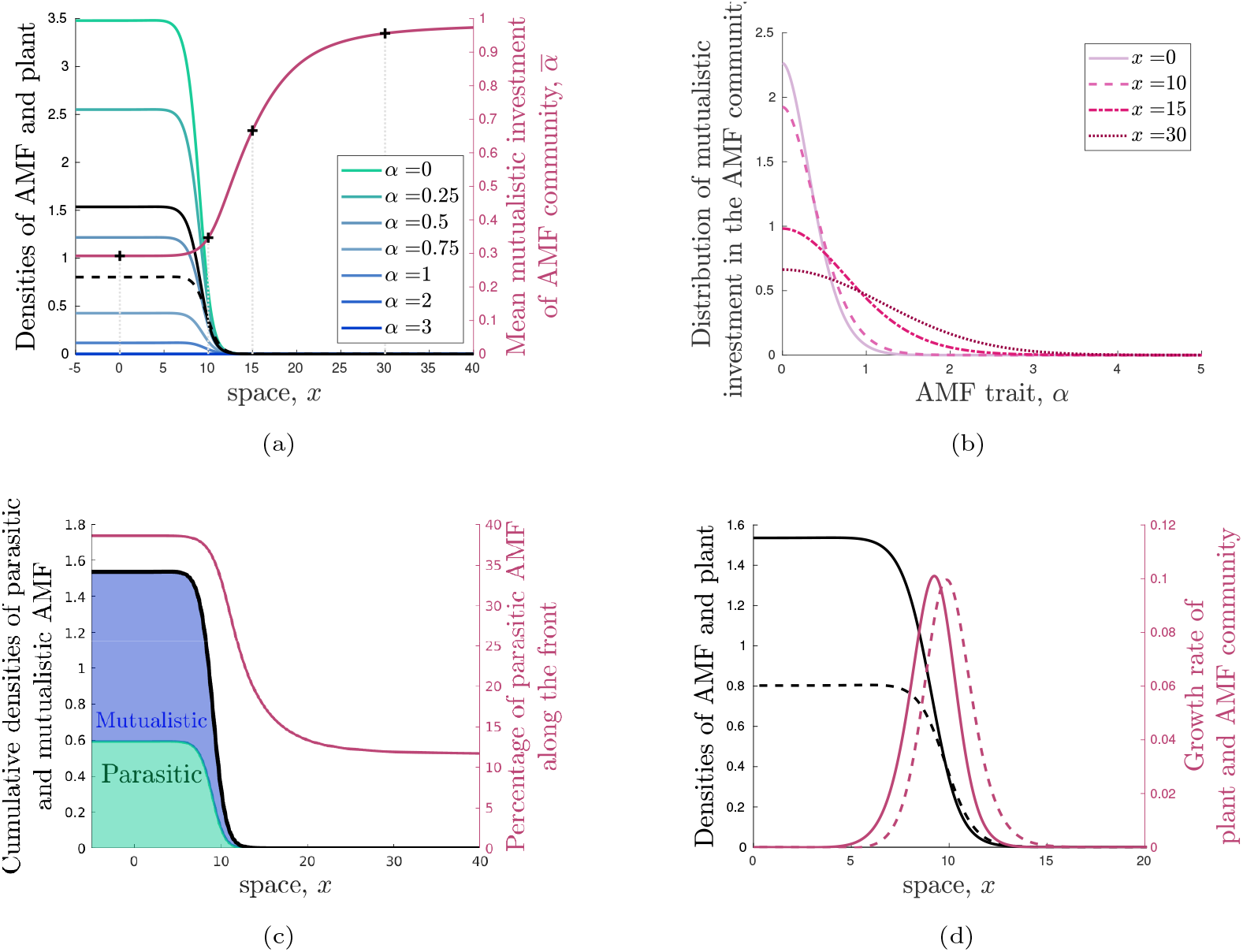
Panel (a), Travelling wave profile. Plot of AMF density for increasing values of *α* (green to blue curves), total AMF density with respect to *α* (black curve), plant density (dashed black curve), and mean AMF trait 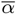 (thick solid red curve). The crosses (and vertical dotted lines) indicate the positions along the travelling wave corresponding to the curves in panel (b). Panel (b), The distribution of the trait *α* at four different positions along the travelling wavefront. For both plots, *β* = 0.4. Panel (c), Cumulative densities of parasitic AMF (*α* ⩽ *α*_*c*_, green area) and mutualistic AMF (*α > α*_*c*_, blue area) inside the traveling front (black curve). Red curve corresponds to the proportion of parasitic AMF along the traveling wave. Panel (d), Growth rate of the plant (dashed red curve) and AMF (solid red curve) as a function of position along the travelling wave (plant - dashed black curve, AMF - solid black curve). Parameter values used for the simulations are: *D*_*m*_ = *D*_*p*_ = 0.1, *d*_*m*_ = 0.01, *Q* = 6, *μ*_*p*_ = *μ*_*m*_ = 0.3, *β* = 0.72, and *α*_min_ = 0 and *α*_max_ = 5.

Note that Fig. 2(a) and Fig. 2(c) are apparently contradictory. The first shows that, in the core of the travelling wave, the parasitic AMF occur at higher density than mutualistic AMF, but that the cumulative density of mutualistic AMF exceeds that of parasitic AMF. This situation arises because there are few values of *α* that qualify as parasitic (*α < α*_*c*_), but many values of *α* that qualify as mutualistic. An examination of Fig. 1(b) confirms this observation: If we consider, for example, the *β* = 0.4 curve and the corresponding gray dotted line indicating *α*_*c*_ (near 0), we can see that the area under the curve to the left of *α* = *α*_*c*_ is indeed small and probably smaller than the area under the curve to the right of *α*_*c*_. This comparison also holds for all of the other curves that appear in Fig. 1(b). Consequently, the cumulative density of mutualistic AMF exceeds that of parasitic AMF, as shown in Fig. 2(c).

#### Fixation probability of parasitic and mutualistic traits

We find that it is evolution among the parasitic symbionts in the population core that gives rise to the mutualistic population at the range edge. Indeed, parasitic symbionts are the most likely common ancestors of mutualistic symbionts (Fig. 3 and Fig. B3). The curve of diamonds in Fig. 3 represents the average fixation probability of symbiont individuals originating from location *x*, that is the probability that their descendants become prevalent at the leading edge of the travelling wave. In other words, this curve shows the probability with which AMF at position *x* along the travelling wave give rise to AMF (which are highly mutualistic) at the leading edge of the wave. Note, however, that at each location *x*, there exists an entire community of symbionts. Consequently, the fixation probability is not uniform among the individuals starting at a given location *x*: the probability truly depends on the value of the trait, *α*, for each symbiont. That is, knowing that an ancestor comes from position *x* is insufficient: We also need to know which of the AMF at that position are the likely ancestors. The diamonds are therefore coloured according to the mean *α* value of the likely ancestors at that position, or the mean mutualistic trait of individuals with respect to this fixation probability. In other words, the colour of each diamond describes the trait value at that location *x* of the ancestors of the symbionts sampled at the leading edge of the front.

**Fig. 3.**
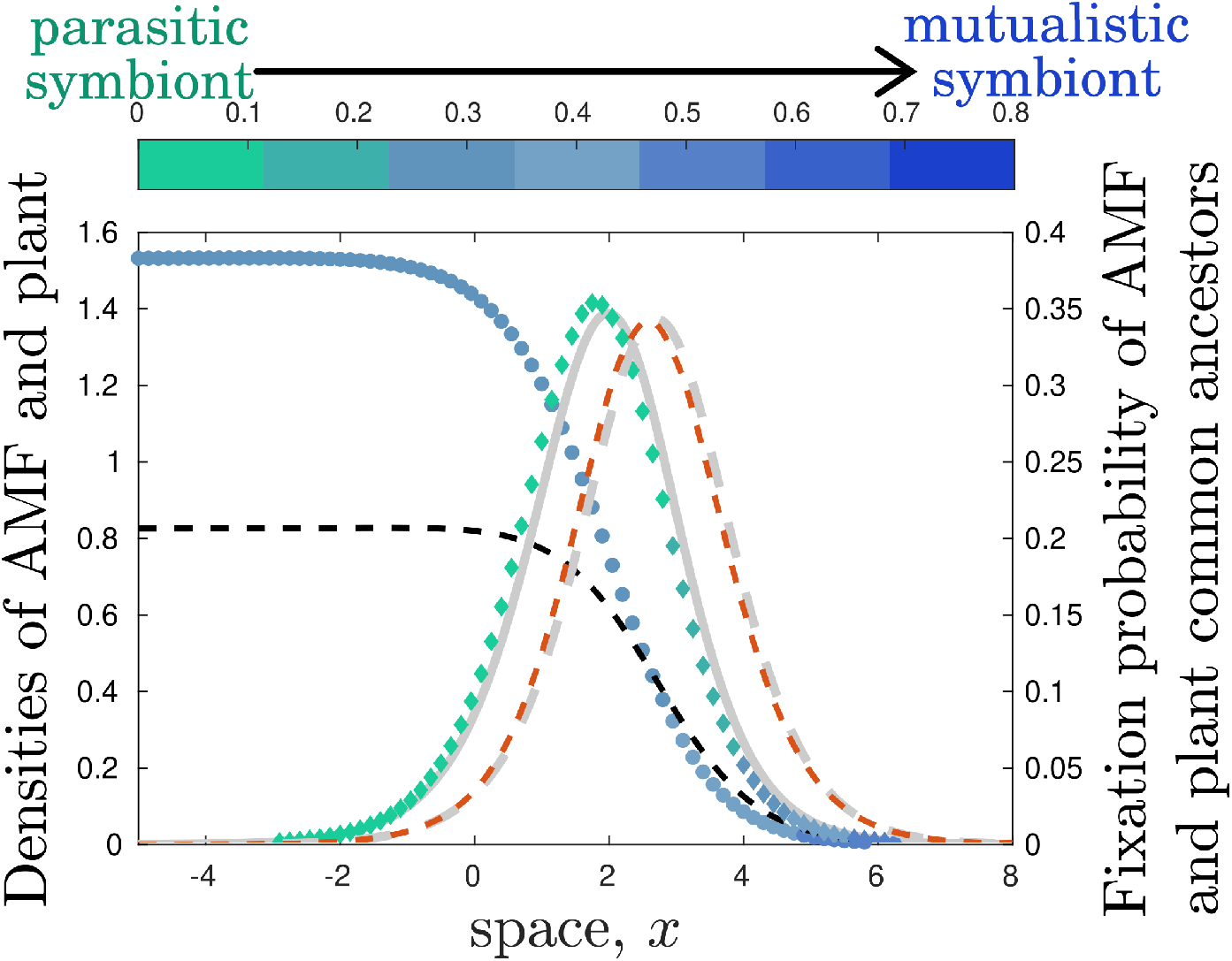
Fixation probability of plant (red dashes) and symbionts (green diamonds) with respect to location in space. Dashes and symbols (dots and diamonds) represent solutions of numerical simulation of Eq. (C65)-(C68) and gray curves are the corresponding approximations obtained from Eq. (6) (C70). We also plot the densities of plants (black dashes) and AMF symbionts (blue dots). The colour of each dot and diamond represents the value of the mean mutualistic trait 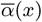 at the corresponding location *x* (where green denotes parasitic and blue denotes mutualistic symbiont communities). Parameter values used for the figure are: *d*_*m*_ = 0.01, *D*_*p*_ = *D*_*m*_ = 0.1, *Q* = 6, *μ*_*p*_ = *μ*_*m*_ = 0.3, *β* = 0.72 and *α*_min_ = 0 and *α*_max_ = 5.

From the curve of diamonds in Fig. 3, we see that: (i) Symbionts just behind the leading edge contribute more to the spread (i.e., the peak of the fixation probability curve is found toward the front of the travelling wave, but behind the leading edge), and (ii) the most likely common ancestors of these symbionts are parasitic symbionts (as shown by the green colour of the peak of the fixation probability curve and, indeed, of most points along this curve).

Thus, the proportion of individuals at the leading edge that come from mutualistic individuals is small (as we can see in the blue right tail of the fixation probability curve), and mutualistic individuals at the leading edge mainly come from parasitic individuals from behind the leading edge of the front.

The mutualistic AMF at the leading edge arise from two sources: (1) offspring of mutualistic AMF at the leading edge, and (2) mutated offspring of dispersed parasitic AMF from behind the leading edge. Since parasites are more abundant than mutualists (see Fig. 2(a)), they make up a larger proportion of the total biomass and thus also disperse more biomass. At the leading edge of the travelling wave, host density is low and selection of AMF at the ecological scale favors mutualism. Thus, of the parasitic AMF that disperse to the leading edge, only those that mutate into mutualists are selected in this region. The mutualistic AMF already at the leading edge also give rise to mutualistic offspring, but given that the biomass of parasitic AMF dispersing to the leading edge far exceeds the biomass of mutualistic AMF already at the leading edge, the proportion of mutualists selected from parasitic ancestors (source (2)) is much larger than the proportion of mutualists arising from mutualistic ancestors (source (1)).

Specifically, the probability of fixation *PF*_*m*_ satisfies the following formula

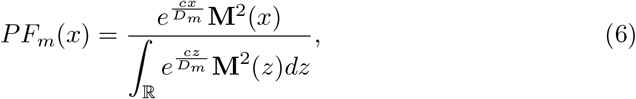

where **M** is the profile of the symbiont density in the moving frame at the spreading speed *c* (see Appendix C.2 for details). The probability of fixation thus depends on the dispersal rate *D*_*m*_, as well as on symbiont growth, through the profile of the symbiont density **M**. We see that the probability of fixation at the leading edge (from mutualistic ancestors) will always be lower than the probability of fixation just behind the leading edge (from parasitic ancestors), meaning that the wave front is mostly composed of parasitic symbionts arriving through dispersal and then mutating into mutualistic symbionts.

Note that the fixation probability curve for the hosts is closer to the leading edge of the travelling wave that of the AMF (Fig. 3). The process, therefore, is that the hosts disperse first and symbionts follow. We emphasize here that, among the symbionts, neither mutualists nor parasites spread faster. The key is that mutualists have an advantage at low host density, while parasites grow larger than mutualists at high host density. Consequently, the mutualists found in regions of low host density (leading edge of the travelling wave) are more likely to stem from mutated parasites dispersing from regions of higher host density (behind the leading edge).

#### Growth rates at the range edges and pushed and pulled waves

A mutualistic community at the leading edge provides a large growth rate to the host plants, leading to a rapid increase in host density (Fig. 2(d)). As plant density increases, the growth rate of the symbionts will also increase, given that the benefit provided by the host to the symbionts increases linearly with host density (see third term in Eq. (1b)). However, it is also true that the strength of selection favoring parasites increases with increasing host density. As the symbiotic community becomes more parasitic behind the wave front, the host growth rate decreases, causing in turn a decrease in the growth rate of the whole symbiotic community. Thus, the maximal symbiont growth rate is located just behind the maximal growth rate of the host population, where host density is large enough to support symbiont growth, but low enough to limit the growth of parasites. This situation is similar to that for the fixation probabilities (cfr. Fig. 3 and Fig. 2(d)). As the host population colonizes first, followed by its associated symbiotic community, the expansion is pulled by the host population, which benefits from the high proportion of mutualists at the range edge, but is pushed by the symbionts, which generate mutualists from mutation within the population core (Fig. 3).

The expansion dynamics of host-associated communities is therefore the result of a unique interaction of pushed and pulled waves. That is, the speed of spread depends on both the characteristics of the wave core as well as its leading edge. We have derived approximations for the speed of spread in SI.C, shown in Fig. 4 (solid curves), which combine information on the mean trait of the population in the core with the growth dynamics at the leading edge of the wave. Our approximations capture the general shape of the speed function, but under-estimate the speed of spread. This difference is partly due to spatial heterogeneity in the mean mutualistic investment of the symbiont community 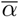, which is not embedded our approximation (i.e., in our analytical approximation 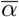 is considered to be constant, while effectively 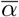 increases in space *x*, see Fig. 2).

**Fig. 4.**
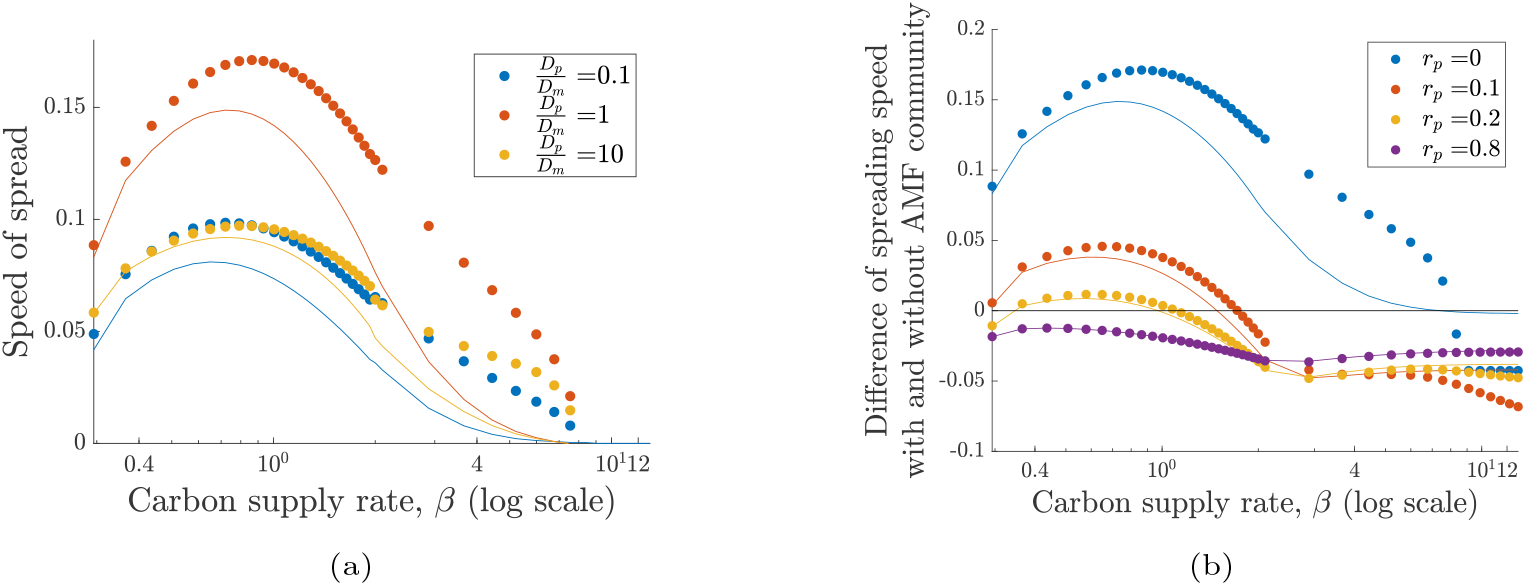
Panel (a), Speed of spread of AMF and plants with different relative dispersal abilities described by the diffusivity ratio (*D*_*p*_*/D*_*m*_): slow plant (blue), slow AMF (orange), and both identical (red); Panel (b) Difference between the speed of spread of a plant with and without an AMF community for various plants with different carbon supply rate (*β*) and different degree of dependence on the symbiont: obligate mutualistic plants (blue, *r*_*p*_ = 0), facultative mutualistic plants with various intrinsic growth rates, *r*_*p*_ = 0.1 (red), *rp* = 0.2 (orange) and *r*_*p*_ = 0.8 (purple). The markers correspond to the travelling wave speed for the model (1) with evolution *d*_*m*_ = 0.01) and the solid curves correspond to the analytical approximation (C59). The parameter values are: *Q* = 6, *μ*_*p*_ = *μ*_*m*_ = 0.3, and *α*_min_ = 0 and *α*_max_ = 5.

### The spread of a host population and its symbionts

Resource supply from the hosts to the symbionts determines the speed of spread of the whole community (Fig. 4). This speed reflects a similar broad unimodal dependence already observed for the plant and AMF densities (cfr. Fig. 1(a) and 4(a)). We find that the speed of spread is maximal when conditions in the core of the population maximise the density of the symbiont community rather than the density of plants (i.e., the maximal speed of spread is obtained around *β* = 0.86, a value close to the one which maximizes symbiont density at equilibrium, *β* = 0.72). Indeed, as discussed in the previous section, the wave of expansion is pushed by the symbionts (Fig. 3 and Fig. 2(d)). Thus, large carbon supply from the host, despite not resulting in maximal plant density, may accelerate range expansion of a given host-symbiont community. Any difference in the dispersal abilities of the host and symbionts decreases the speed of spread (compare the blue and yellow curves with the red curve of Fig. 4(a)), although the carbon supply rate that maximizes the speed of spread remains essentially the same (compare the location of the peak in the blue, yellow, and red curves of Fig. 4(a)).

As the wave of expansion is pulled by the plants (Fig. 3 and Fig. 2(d)), the speed of spread increases if the plants can grow in the absence of the symbiont community (Fig. C4(b)). However, the speed advantage provided by the symbiont community depends on the obligate degree of the plant. When considering the difference between the speed of spread in the absence and presence of symbionts, we show that symbionts can induce either a positive or negative effect on the speed of spread (Fig. 4(b)). When the plants have a small intrinsic growth rate and need the symbiont community for growth, symbionts enhance the speed of spread. For plants with a large intrinsic growth rate, however, the presence of symbionts reduces the speed of spread. The speed load induced by the symbionts depends on the cost of symbionts to the host, which depends on the rate of carbon supply *β*. In conclusion, strategies that decrease the dependency of hosts on their symbionts (such as a transition from obligate to facultative mutualism) or, alternatively, strategies that increase symbiont density in the core of the population (such as selection for a carbon supply rate that maximizes symbiont density), enhance the speed of spread of the whole host-symbiont community.

## Discussion

### The Eco-evolutionary dynamics of host-associated communities

There have been several calls for investigation of the dynamics that emerge when evolutionary and ecological changes in symbiotic communities occur at the same timescale (Koskella et al, 2017; Fitzpatrick et al, 2020; Drew et al, 2021b). Evolutionary and ecological forces are known to drive the spread of pathogens in host populations (Restif, 2009; Day et al, 2020), but our understanding of eco-evolutionary dynamics involving mutualistic symbionts remains limited. Here we present a theoretical framework that allows us to investigate, under minimal assumptions, the linked population dynamics, spread, and evolution of a host population and its associated mutualistic and parasitic symbionts. We show how mutualism can emerge from a parasitic community of symbionts, and stably persist in a community in which parasites and mutualists coexist and rapidly evolve along the parasitism-mutualism continuum (Lin and Koskella, 2015; Rogalski et al, 2021). Our framework accounts for unique characteristics of host-associated communities, such as their short generation time and horizontal modes of transmission, and constitutes the basis for further investigation of the eco-evolutionary dynamics of host-symbiont communities.

In our model, coexistence of parasites and mutualists occurs thanks to two key features: (1) a mutation-selection balance acting on the symbionts at the individual scale, and (2) a positive eco-evolutionary feedback acting at the host-community level. At the individual scale, parasitic symbionts benefit from a higher fitness than mutualistic symbionts, because they receive the same benefit from the host at a smaller cost. However, the average mutualistic investment of the whole community, and not the individual contribution of each symbiont, determines whether a community will establish and be mutualistic as a whole, as observed in previous modeling work (Archetti and Scheuring, 2011, 2013; Martignoni et al, 2020). Thus, in our context, the evolution of mutualism looks similar to the evolution of altruism. Even though the emergence of altruism is a fraternal transition (i.e., arises from a division of labour among individuals of the same species), whereas the transition from parasitism to mutualism is egalitarian (i.e., results from the association of different species to complement their function), both result from the conflict between two levels of selection: the individual-level, favouring parasites, and the group-level, favoring altruists or mutualists (Wilson and Sober, 1989; Van Baalen and Rand, 1998; Simon et al, 2013).

Previous work has shown a strong shift from mutualism to parasitism with increasing availability of resources, with more beneficial symbionts dominating the community when host productivity is low, and parasites dominating when host productivity is high (Schwartz and Hoeksema, 1998; Hochberg et al, 2000; Hochberg and van Baalen, 2000; Neuhauser and Fargione, 2004). Similarly, in our simulations the selective advantage of parasites increases with the amount of resource available (i.e., with host density). Thus, high host density promotes the establishment of parasitic communities and, conversely, low host density reduces the selective advantage of parasitic symbionts, causing the community to become more mutualistic on average. Fluctuations in selection pressures due to, for instance, nonlinear public goods Archetti and Scheuring (2013), genotype-environment interactions (Parker, 1995), negative frequency-dependent selection (Bever, 1999; Brown and Tellier, 2011) or selection mosaics (Thompson, 2005) are known to maintain variation in partner quality (Mitchell-Olds et al, 2007). Here we show that fluctuation in selection pressures, due to variation in host density, can induce a shift from parasitism to mutualism in a symbiotic community.

Interesting future directions could consider how the inclusion of additional mechanisms, such as host control and different symbiont transmission modes, may improve the quality of the mutualistic interactions between hosts and symbionts. For example, host phenotypic plasticity in its mutualistic investment toward the symbionts may facilitate the evolution of more mutualistic traits (Koskella and Bergelson, 2020; Hou et al, 2021), even though the evolution of mutualism among hosts occurs after the emergence of mutualism among symbionts (Ledru et al, 2022). Strict vertical transmission of symbionts could also enhance evolution toward stronger mutualistic communities (Ewald, 1987; Sachs et al, 2004). Competition between hosts can also shape community structure: More mutualistic symbiotic communities should provide a fitness advantage to their host, which could drive the selection for communities with a lower proportion of parasites (Hartnett et al, 1993; Jones et al, 2012). Extension of our current model would allow us to investigate the evolutionary potential of these additional mechanisms.

### Community structure at the range edges

Range expansion can result from various mechanisms involving growth and dispersal. Generally, we distinguish between two types of expansions: those that are pushed *versus* those that are pulled. In a pulled propagating wave, the population expands its range thanks to a higher growth rate at the population edges, while in a pushed wave population expansion is driven by a higher growth rate in the core (but see (Gandhi et al, 2016; Miller et al, 2020; Erm and Phillips, 2020)). Here, host-symbiont communities follow a combined expansion dynamic in which the wave of expansion is pulled by the hosts and pushed by the symbionts.

When host-symbiont communities expand their range, the selective pressure on symbionts becomes a function of space, due to changes in host density along the travelling wave, with strong selection favouring parasitic symbionts in the population core (where host density is large) but not at the population edges. The low density of plants at the population edges promotes the formation of a more mutualistic community, causing plant growth rate to be maximal at the wave front. Thus, the host plants pull the expansion wave by providing new resources to their obligate symbionts. In contrast to the pulling dynamics of the plants, the wave of expansion is pushed by the symbionts from the population core. The symbiotic community in the population core is more parasitic than that at the wave front, due to higher resource availability. As discussed in the previous section, parasitic symbionts are the common ancestors of good mutualists at the wave front. Thus, these symbionts are pushed from the population core into the front and closely follow their hosts in their expansion dynamics.

Spatial self-organization has been found to increase the abundance of mutualistic symbionts during range expansion in theoretical models (Momeni et al, 2013; Van Dyken et al, 2013) and synthetic microbial communities (Pande et al, 2016; Amor et al, 2017; Rodríguez Amor and Dal Bello, 2019), with cheaters or parasitic symbionts lagging behind the mutualists at the leading edge. Emerging evidence also indicates that under stressful conditions plants may actively ‘cry for help’ (i.e., by secreting chemical compounds) and recruit beneficial symbionts (Schuman et al, 2015; Rizaludin et al, 2021). In our work, spatial selection occurs in the absence of built-in mechanisms. Instead, it is spontaneously driven by lower host density at the leading edges, which reduces the selection strength against mutualists in this region.

Invasion by alien organisms is often studied from the perspective of a single population, despite the fact that the success of an invasion might require the simultaneous successful spread of the focal population and its invisible symbionts (Dickie et al, 2017). Eradication and restoration strategies should therefore consider both hosts and symbionts as active and tightly-linked players in invasion dynamics. As conservation strategies to limit pulled and pushed invasions differ (Gandhi et al, 2016), the combined push-pull dynamics of linked host-symbiont invasions may require novel containment strategies (Taylor and Hastings, 2004). For example, limiting the invasive spread of species expanding as a pulled wave is currently accomplished by eradicating invaders at the population edges. However, this strategy may be inefficient if expansion and evolution of the symbionts in the population core facilitates the quick restoration of the population at the range edges. Our general framework may be adapted to host-symbiont populations expanding over a heterogeneous landscape, such as a fragmented or disturbed habitat (Willing et al, 2021), and to consider range overlaps with native populations (Dickie et al, 2017).

### The spread of a host population and its symbionts

The effect of mutualistic interactions between hosts and symbionts on their range expansion is unclear in the existing literature. On the one hand, the growth benefits provided by mutualism can increase the speed of spread of the host, while on the other hand the absence of mutualistic partners at the range edge may produce an Allee effect that limits the rate of spread into new territories (Stanton-Geddes and Anderson, 2011; Kubisch et al, 2014; Fowler et al, 2023). Our framework allows us to disentangle these two tendencies and study how resource exchange (determining fitness of hosts and symbionts along the expansion range) and differences in host and symbiont dispersal abilities can affect the speed of spread of the community.

The ‘enemy release hypothesis’ postulates that species can escape their enemies (such as predators, pathogens, parasites, or herbivores) when introduced into a new range (Torchin et al, 2003; Colautti et al, 2004). A similar effect can occur at the edge of an expanding population, if enemies disperse more slowly than their hosts (Fagan et al, 2002; Nomikou et al, 2003). In contrast, if the linked species is beneficial, an expanding host population may also escape its mutualistic symbionts (Dickie et al, 2017; Shaw, 2022), to the detriment of its speed of spread. A similar behaviour occurs in cooperative systems, where the phenotype with the slowest speed of spread limits the spreading speed of the community Li et al (2005).

In agreement with these results, we found that differences in the dispersal abilities of hosts and symbionts may reduce expansion rates. Thus, symbiont inoculation at the leading edges can also contribute to accelerating range expansion if symbionts have limited dispersal ability. We also found that the optimal carbon supply rate, i.e., the one that maximizes AMF density at equilibrium, corresponds to the rate that maximizes the speed of spread of the whole host-symbiont community. Thus, host investment in the mutualistic relationship with symbionts may be considered an evolutionary strategy to increase the colonization speed of the host-symbiont community, and might also compensate for an eventual decrease in the speed of spread due to a lack of symbionts at the leading edge.

Our results also show that the speed of spread of host-symbiont communities can be maximized with reduced host dependence on the mutualism (from obligate to facultative). It follows that the evolution of reduced dependence of a host on its symbionts at the range edges of an expanding population can potentially facilitate plant invasion (Seifert et al, 2009). Note, however, that we only included one axis of benefit to the hosts, that is, one symbiont-supplied resource. Mutualistic interactions can be multidimensional; AMF, in particular, have been found to provide a suite of benefits to their hosts, such as protection against pathogens or abiotic factors (Smith and Read, 2010; Fowler et al, 2023). The inclusion of these other benefit axes might reduce the benefit of a host reducing its dependence on symbionts.

Our model could be extended to consider the evolution of obligate mutualism from facultative mutualism. Indeed, hosts that are facultative mutualists may reach higher densities in the absence of symbionts. Subsequent colonization of these hosts with parasites, and the evolution of some mutualistic traits in these parasites, may be sufficient to drive a transition from parasitism to mutualism in the community. Future studies are needed to consider how, once mutualism has emerged, host adaptation may lead to increased investment into the mutualistic relationship with symbionts, rather than into self-growth (Ledru et al, 2022). Similarly, the model could be extended to consider the evolution of microorganisms from free-living to host-associated states (Drew et al, 2021b).

Finally, it might be interesting to investigate dynamics of co-evolution between the mutualistic investment of symbionts and their dispersal ability. Indeed, in some circumstances dispersal ability has been found to evolve during range expansion (Urban et al, 2007; Brown and Vellend, 2014). A recent study (Ledru et al, 2022) has shown that co-evolution between mutualistic effort and dispersal results in mutualistic symbionts that disperse locally (i.e., have low dispersal ability), while parasitic symbionts disperse over longer distances (i.e., have high dispersal ability). Such negative correlation between dispersal ability and mutualistic quality has also been observed in the context of altruism (Koella, 2000; Le Galliard et al, 2005; Hochberg et al, 2008; Eldakar et al, 2010; Purcell et al, 2012; Mullon et al, 2018), and between local interactions and evolution of avirulence (Boots and Mealor, 2007). In a range expansion context, this correlation might be reversed because mutualistic symbionts have an evolutionary advantage at the leading edge.

### Conclusion and future work

Current ecological and evolutionary theory fails to explain the complex dynamics structuring stable host-symbiont communities (Koskella et al, 2017). This gap is at least partly due to unique characteristics of these communities, such as rapid evolution, interdependent fitness, and horizontal transmission of symbionts, which make study and analysis challenging. Here we present a theoretical framework that contributes to our understanding of host-symbiont linked population dynamics, evolution, and spread, with possible implications for microbiome research (Koskella et al, 2017; Fitzpatrick et al, 2020) and for the management of linked plant-microbial invasions (Dickie et al, 2017).

## Acknowledgments

MMM acknowledges the Azrieli Foundation. MMM and OK acknowledge the Israel Science Foundation (ISF) (grant number 1826/20), the Gordon and Betty Moore Foundation, and the United States-Israel Binational Science Foundation (BSF). RCT acknowledges the Natural Sciences and Engineering Research Council (NSERC) of Canada Discovery Grants Program, grant number RGPIN-2022-03589, and the University of British Columbia Okanagan Institute for Biodiversity, Resilience, and Ecosystem Services. JG acknowledges ModEcoEvo project funded by the Université Savoie Mont-Blanc.

## Statements and declarations

- The authors have no competing interests to declare that are relevant to the content of this article.
- All authors certify that they have no affiliations with or involvement in any organization or entity with any financial interest or non-financial interest in the subject matter or materials discussed in this manuscript.
- The authors have no financial or proprietary interests in any material discussed in this article.
- All authors agree with the present manuscript
- For their simulations, authors used Matlab R2022. All the codes developed for the manuscript are available on GitHub https://github.com/garnieji/AMF_Evolution

### Authors’ contributions

MMM: Conceptualization, Methodology, Investigation, Writing - Original Draft, Writing - Review & Editing, Visualization. RT: Conceptualization, Methodology, Investigation, Data curation, Writing - Review & Editing, Visualization, Funding acquisition. OK: Writing - Review & Editing, Funding acquisition. JG: Conceptualization, Methodology, Software, Formal analysis, Investigation, Data curation, Writing - Review & Editing, Visualization, Funding acquisition.

## Appendix A No evolution no space, competitive exclusion and parasitic threshold

We first look at the dynamical system without spacial component and evolution of the trait *α*, and we only focus on obligate hosts, for which the intrinsic growth rate *r*_*p*_ = 0).

### Monomorphic population of AMF

We first look at a population of symbionts that are monomorphic, that is all the individuals share the same trait *α*. In this situation, the system is reduce to the following differential equations for the biomass of plant *P* and the biomass of the monomorphic symbionts *M* :

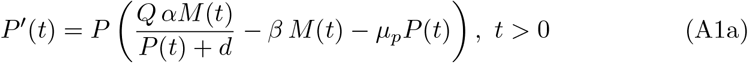

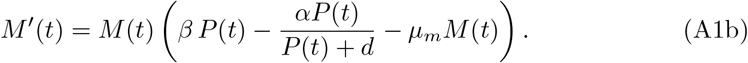

From the analysis of Martignoni et al (2020), the system admits the extinction steady state (0, 0), which might be stable if *d < α/β* and a stable positive steady state (*P, M*) if the following properties are satisfied:

- *Q >* 1
- *α* ⩾ *α*_*c*_(*β*) where *α*_*c*_(*β*) satisfies

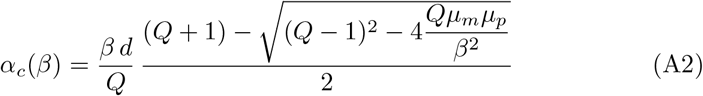
- *μ*_*p*_*μ*_*m*_ ⩽ *β*^2^(*Q* − 1)^2^*/*(4*Q*).

The assumption on *α* shows that the mutualistic investment of the symbiont needs to be large enough to persist with the plant, that is *α* ⩾ *α*_*c*_ where the threshold satisfies

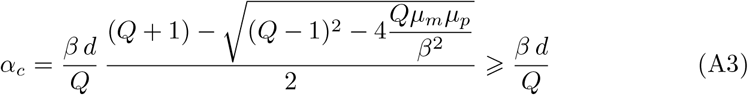

which means that the symbiont provides a positive benefit to the plant when the plant is at low density. In addition, the analysis first points out that if the monomorphic population of AMF have a low mutualistic quality *α* below the threshold *α*_*c*_, then the system goes extinct. As such, we say the a symbionts is *parasitic* if its mutualistic investment *α* is below *α*_*c*_, while it is *mutualistic* if *α > α*_*c*_.

### Polymorphic population of AMF and competitive exclusion

On the other hand, if the symbiont population is initially polymorphic, with an initial trait density *m*(*t, α*) with *α* ∈ [*α*_min_, *α*_max_], then the population will converges toward a monomorphic population with mutualistic quality *α*_min_. If this quality is below the critical value *α*_*c*_, the population will go extinct.

The population densities (*P, m*) satisfies in this case the following model:

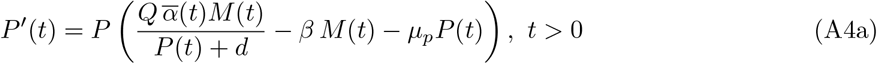

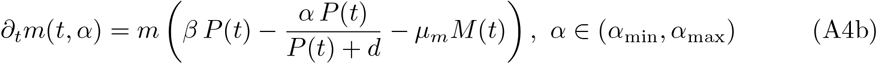

where the quantity 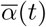 is the mean AMF trait in the community at location *x* and *M* (*t*) is the total biomass of AMF, defined by

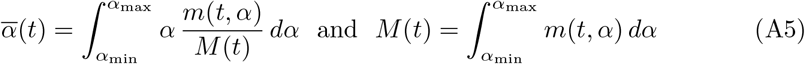

As the result, the total biomass of AMF satisfies the following equation

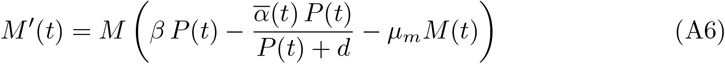

and the trait distribution *ϕ*_*m*_(*t, α*) = *m*(*t, α*)*/M* (*t*) is described by the following model

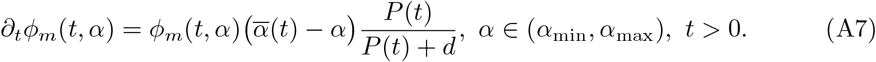

From the definition of the mean trait 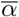, it satisfies 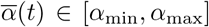 for all time *t >* 0 and for all 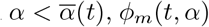is decreasing over time, while *ϕ*_*m*_(*t, α*) is increasing for 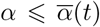. Consequently, 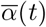 is decreasing over time, and it converges to *α*_min_, while *ϕ*_*m*_ converges toward the dirac mass at *α* = *α*_min_.

If the minimal value of the trait *α*_min_ is below the persistence threshold *α*_*c*_ then the population goes extinct. So in absence of diversification of trait *α*, the population experience a competitive exclusion that can expose the population to extinction.

## Appendix B Evolution, no dispersal, analysis

To understand the co-evolution of a plant with its AMF community, we first look at the diversification of AMF for a given plant at a fixed location. In this situation, the plant with biomass *P* (*t*) is able to provide carbon for the AMF with trait *α* and density *m*(*t, α*), at a rate *β*.

The model (1) then becomes

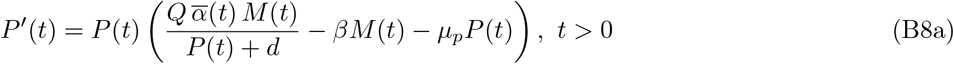

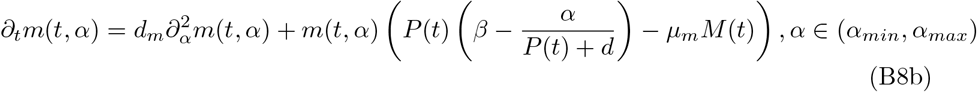

We here aim to understand the AMF community that emerges from the interaction with the plant and the evolutionary process. More precisely, we aim to characterize the steady states of this model (B8). We first prove the existence and some properties of the non trivial steady state and then we provide an approximation which corresponds to special case where *α*_max_ = ∞.

Let us first write the model (B8) with different variables. First, if we integrate over *α* the equation for *m*, we obtain the following system of equations for the total biomass of AMF *M* and plant *P* :

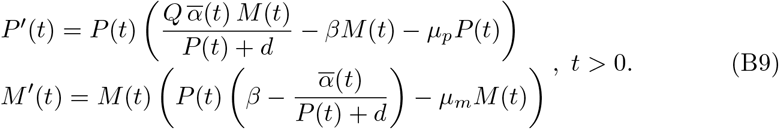

Then, let us write an equation for the trait distribution of AMF *ϕ*_*m*_(*t, α*) = *m*(*t, α*)*/M* (*t*). Using equation for *m* and *M*, we obtain

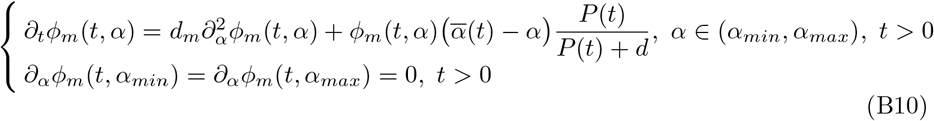

### B.1 Steady state

A non trivial steady state (*P, m*(*α*)) of the model (1) without spatial dispersal (*D*_*p*_ = *D*_*m*_ = 0, should satisfy the two following problems. First, the biomass of the plant *P* and the total biomass of AMF *M* = ∫ *m*(*α*)*dα* solve the following system

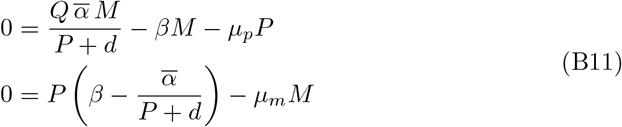

where 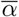 is the mean trait of the AMF distribution at steady state. And the trait distribution at equilibrium *ϕ*_*m*_(*α*) = *m*(*α*)*/M* satisfies the following elliptic problem

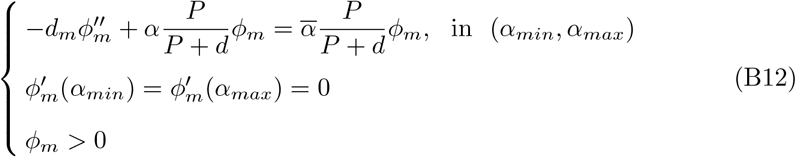

#### Proposition 1.

*If, the diversification rate d*_*m*_ *>* 0 *and the following inequality holds true*

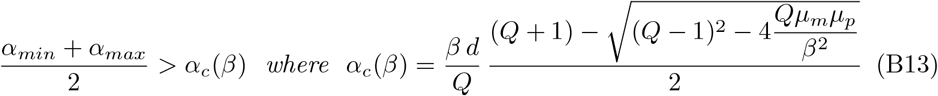

*then, there exists a steady state of the model* (B8).

*In addition, the mean trait* 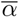*at equilibrium satisfies the following property*

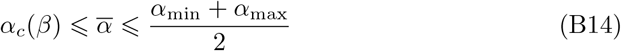

*Proof of Proposition 1*. If a non trivial steady state (*P, m*(*α*)) of the system (B8) exists then the total biomass (*P, M*) should satisfy the following system, which depends on 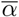,

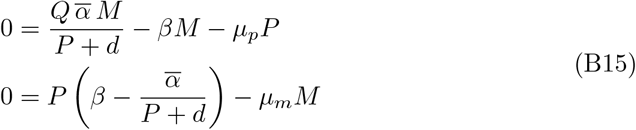

From Martignoni et al (2020), we know that the system admits at least a positive solution, if the following conditions holds true

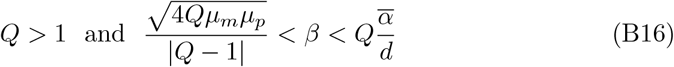

Under this condition, the following equilibrium exists

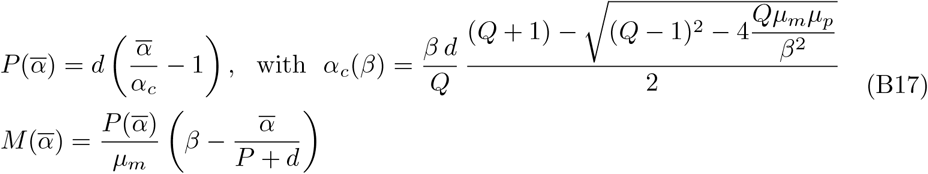

In addition, we can show from (B10) that 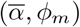 solves the following problem, which depends on *P*

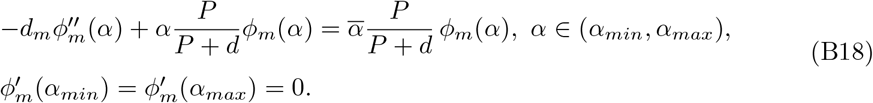

However, we know that for any positive *P*, the Neumann principal eigenproblem

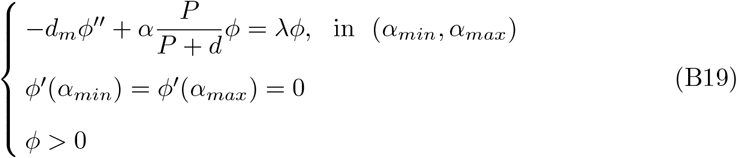

admits a principal eigenpair (*λ*_*P*_, *ϕ*_*P*_), normalized by 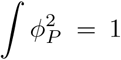 and we have the variational formula

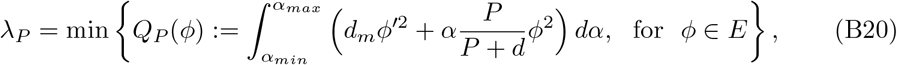

where

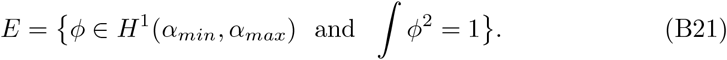

If the parameter *d*_*m*_ *>* 0, the eigenpair (*λ*_*P*_, *ϕ*_*P*_) satisfies the following properties

i. for all *P >* 0, we have

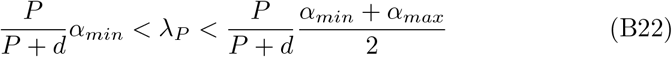
ii. the function *P* ↦ *λ*_*P*_ is increasing and concave in (0, ∞), and

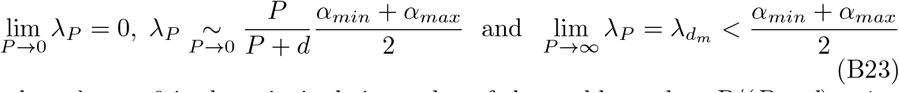

where 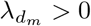 is the principal eigenvalue of the problem when *P/*(*P* + *d*) = 1.
iii. the eigenfunction *ϕ* is is decreasing on [*α*_min_, *α*_max_] and it is strictly concave on [*α*_min_, *λ*_*P*_) and strictly convex on (*λ*_*P*_, *α*_max_].

Since 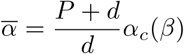 from (B17), we know that there exists a unique positive *P* such that 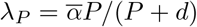 if and only if the following inequality holds true

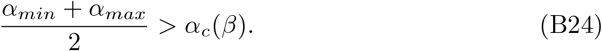

This inequality ensures the existence of a unique steady state 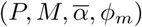 which satisfies simultaneously (B17) and (B12).

Moreover, this inequality shows that when *β* → *β*_*max*_ such that inequality (B24) is an equality, then the mean trait in the AMF population converges towards 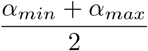 and thus the distribution *ϕ*_m_ converges towards the uniform distribution *ϕ*_*m*_(*α*) = 1*/*(*α*_*max*_ − *α*_*min*_) for all *α* ∈ (*α*_*min*_, *α*_*max*_).

So, if *β* = *β*_*max*_ and competition between AMF all sharing one plant, we expect a uniform distribution of AMF biomass across the range of *α* values (Fig. 1b).

In addition, if the plant density is low *P* → 0 then the mean AMF trait should be close to 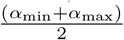 and thus the AMF distribution should be uniform over *α*. As a result, when the plant density is low, we expect the AMF distribution to be uniform

Moreover, the presence of mutation through the diffusion operator is crucial for the existence of a non trivial positive steady state. In absence of mutation, the competitive exclusion drives the population toward extinction.

### B.2 Approximation of the steady states

We aim to find some approximations for 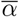, *P, M*, and *m*. First, we assume that

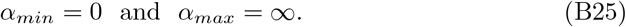

#### Exact solution for the eigenpair problem

In this situation, we can compute an explicit solution of (B19) using the Airy function Ai, which satisfies Ai^*′′*^(*z*) − *z* Ai(*z*) = 0 for all *z* ∈ ℝ. Indeed, the following function *ϕ* is a positive solution of (B19):

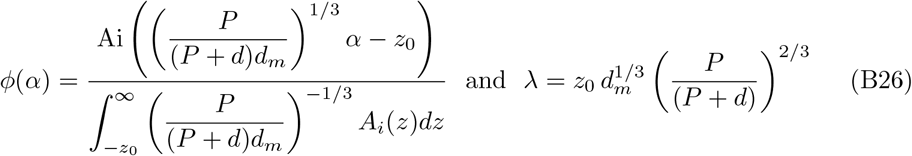

where −*z*_0_ is the maximal value such that Ai achieves a maximum at −*z*_0_ and the mean trait 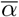 and *P* solves the following system:

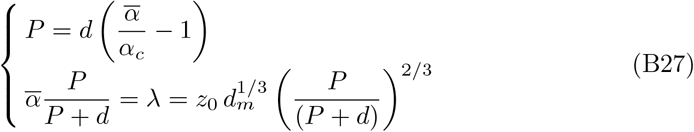

The following system can be reduced to 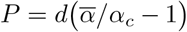 and the mean trait 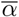 is the real positive root of the following polynomial function 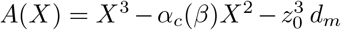.

### B.3 Effect of the diversification rate *d*_*m*_ and plant biomass

We can write the previous system (B27) in the following form

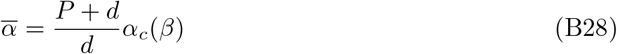

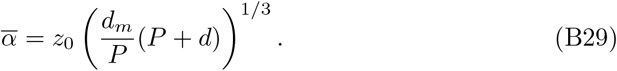

First, the mean trait 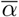 is increasing with the diversification rate *d*_*m*_. As a result, the biomass of the plant as well as the total biomass of AMF is also increasing with the diversification rate *d*_*m*_ (Fig. B1. Secondly, the plant biomass *P* has antagonistic effects on the mean trait 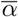. The ecological dynamics, described by (B28), increases the mean trait with the biomass of the plant *P* . A more mutualsitic community will support a larger plant (see Martignoni et al, 2020). While, the evolutionary dynamics described by (B29), decreases the mean trait when the bimoass of the plant increases. Indeed, in the evolutionary dynamics, the strength of selection, tha is *P/*(*P* + *d*), is driven by the biomass of the plant. If the biomass of the plant is large, selection is strong and it drives the trait distribution toward small values of *α*. Conversely, when the biomass of the plant is small, selection is weak and the mutations drive the trait distribution toward larger *α*. The equilibrium results from the balance between the ecological forces which promotes mutualism and evolutionary forces enhancing parasitism.

### B.4 Proportion of parasitic symbionts

In previous section, we introduce the critical value *α*_*c*_ of the trait, such that if the plant is in association with a single symbiont with trait *α* ⩽ *α*_*c*_, then the plant-symbiont system goes extinct. Conversely, the system survives if *α > α*_*c*_. The critical value *α*_*c*_(*β*) does depend on the mutualistic investment of the plant *β*.

In a polymorphic community, where the trait can range from *α*_min_ to *α*_max_ such that *α*_min_ *< α*_*c*_ *< α*_max_, the symbionts can be split in two groups: parasitic symbionts (*α < α*_*c*_) and mutualistic symbionts (*α* ⩾ *α*_*c*_). We investigate the proportion of parasitic and mutualistic symbiont in the community for different plant (Fig. B2). The proportion is defined by

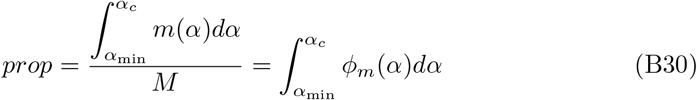

With the approximation of *ϕ*_*m*_ defined in (B19), we obtain

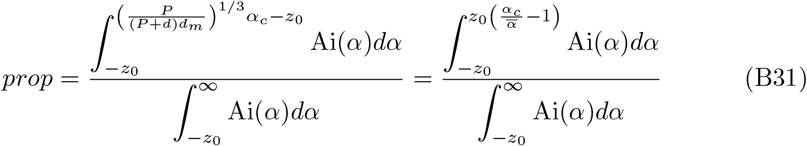

**Fig. B1.**
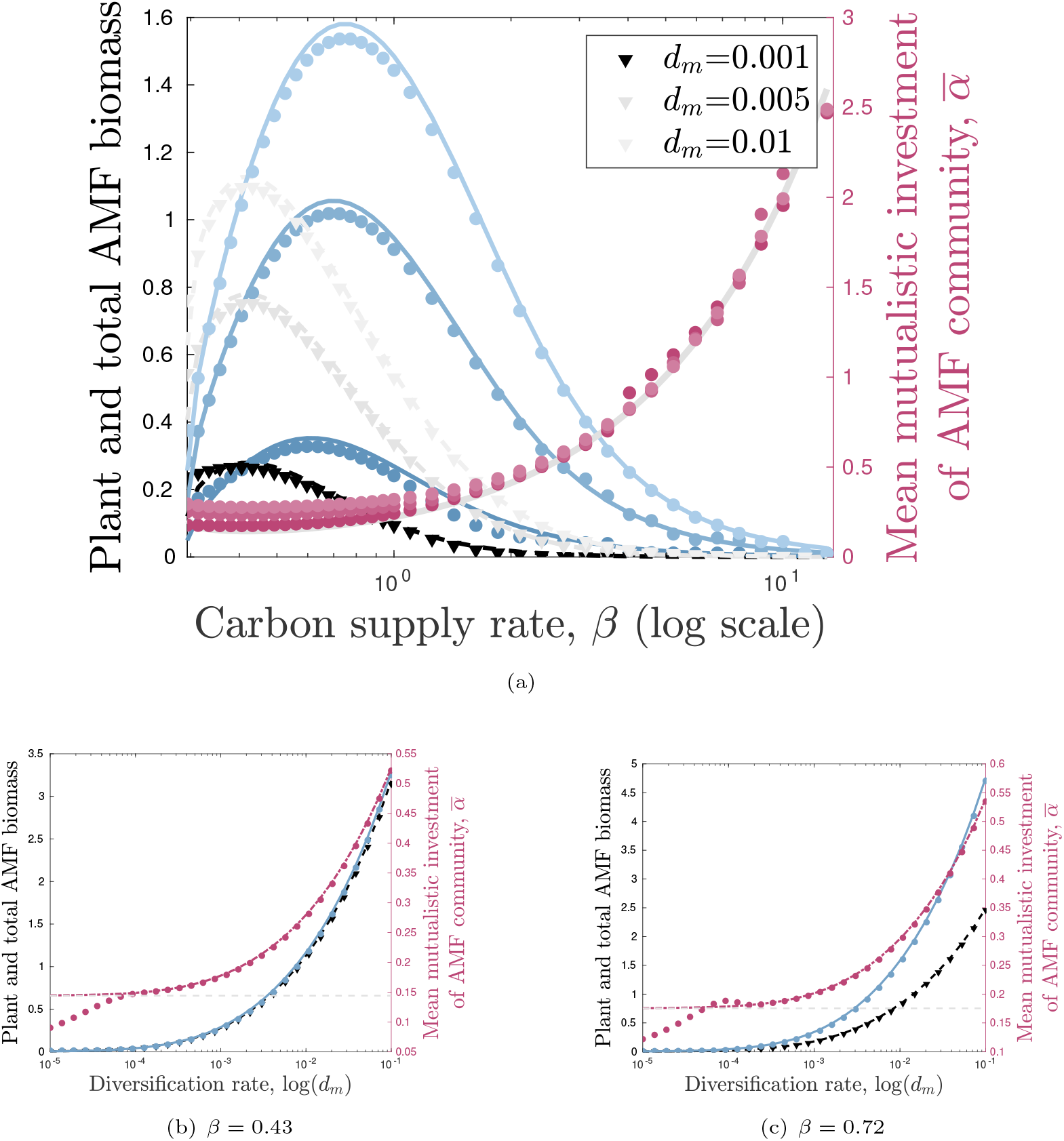
Effect of the diversification rate *d*_*m*_ on the biomass of the plant (black curves and triangles) and the total biomass of the AMF (blue curves and dots), and the mean mutualistic investment 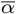 (red curves and dots) at eco-evolutionary equilibrium. For the simulations, the parameters are: *Q* = 6, *μ*_*p*_ = *μ*_*m*_ = 0.3, and *α*_min_ = 0 and *α*_max_ = 5.

On the one hand the proportion of parasitic symbionts increases with the plant biomass if the parasitic threshold *α*_*c*_ remains constant. And the proportion is decreasing with the diversification rate. On the other hand, the proportion of parasitic symbionts depends on the ratio between the critical threshold *α*_*c*_(*β*) and the mean trait 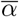. Since 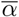 is increasing with respect to the diversification rate *d*_*m*_, the proportion of parasits in the community decreases with the diversification rate, as expected by the previous results (Fig. B2(a-b)).

At equilibrium, the expression of *α*_*c*_ and the relationship between *P* and 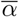 (B28) provide the following expression:

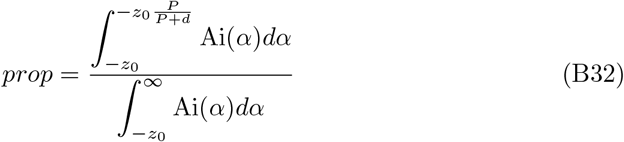

In this case, the porportion of parasitic symbionts depends on the bimoass of the plant at equilibrium. The effect of the mutualistic investment of the plant on the proportion is thus not monotonic because, the biomass of the plant is not monotonic with the parameter *β*. However, we have shown that the plant biomass at equilibrium is increasing and then decreasing with respect to *β*. The proportion of parasitic symbionts is decreasing and then increasing with respect to the mutualistic investment of the plant *β* (Fig. B2).

### B.5 Lineages among the symbionts community

We aim to understand the lineages of individuals present at time *t* in the community at equilibrium. We first track the offspring of individuals, which are present initially, using the inside dynamics approach developed by Roques et al (2012). Individuals are labeled, and transmit their label to their offspring. Since individuals only differ by their label and their trait, each label *k* ∈ ℕ corresponds to a neutral fraction of density *υ*^*k*^ inside the community at equilibrium *m*(*α*). Initially we assume that

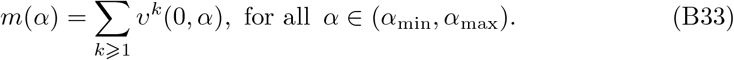

Their dynamics is described by

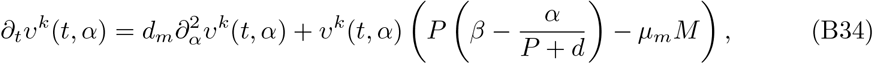

where 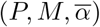 are the equilibrium satisfying (B11)-(B12) and the Neumann boundary conditions

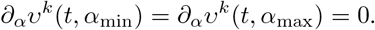

The linear operator associated to this equation is compact and self-adjoint. Moreover, we know that *m*(*α*) is the eigenvector associated to the principal eigenvalue 0. Then from classical semi-group theory (Henry, 1981; Pazy, 1983), the solution *υ*^*k*^ converges uniformly in space toward the following quantity 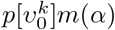 where the positive scalar *p*[*υ*_0_] is defined by

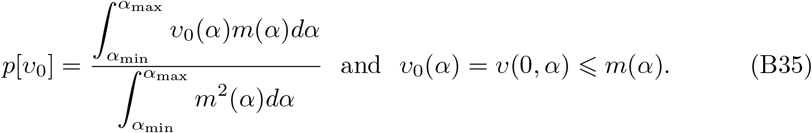

**Fig. B2.**
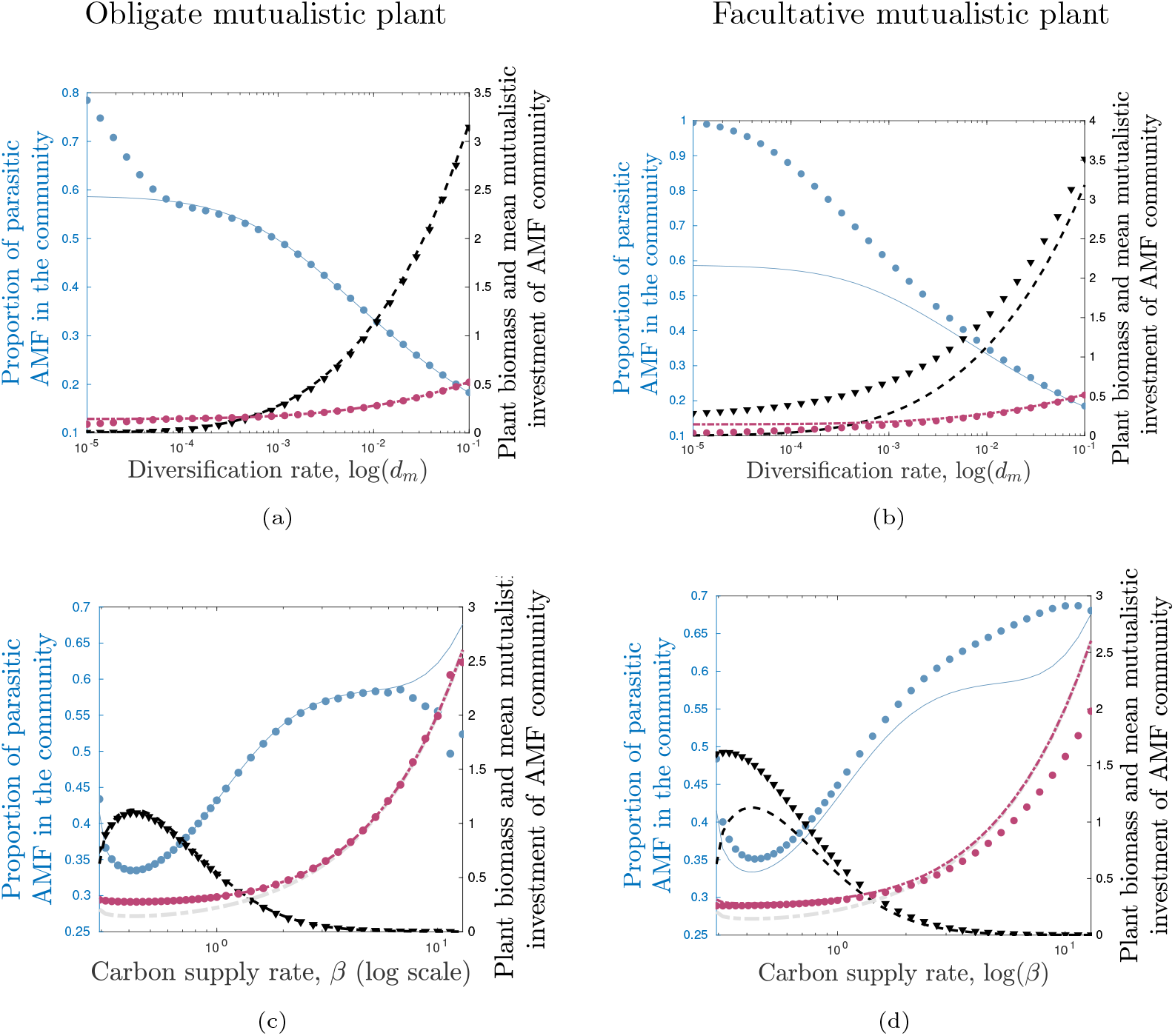
Proportion of parasitic symbionts in the symbionts community associated with an obligate (panel (a-c) *r*_*p*_ = 0) or facultative (panel (b-d) *r*_*p*_ = 0.2) mutualistic plant with various carbon supply ability *β*. Blue dots corresponds to the proportion at equilibrium from the model (1) and the blue curve corresponds to the approximation defined by (B26). Biomass of the plant (black curve and triangles) and mean mutualistic investment of AMF in the community (red curve and dots). Grey curve corresponds to the mutualistic threshold *α*_*c*_. Dashed black line in panel (d) is the approximation for the obligate mutualistic plant biomass. For the simulations, the parameters are: *d*_*m*_ = 0.01, *Q* = 6, *μ*_*p*_ = *μ*_*m*_ = 0.3, *β* = 0.43 and *α*_min_ = 0 and *α*_max_ = 5.

Using this forward approach, we are able to define a backward *ancestral process Y*_*s*_, for each individual with trait *α* at time *t* (see Forien et al (2022) for more details on the derivation). More precisely, the ancestral process at time *s* aims to describe the trait of the ancestor (alive at time *t* − *s*) of an individual sampled randomly among individuals with trait *α* at time *t*. The ancestral process *Y*_*s*_ is related to the previous model through the following relationship for all (*t, α*) ∈ (0, ∞) × (*α*_min_, *α*_max_) and *k* ∈ ℕ:

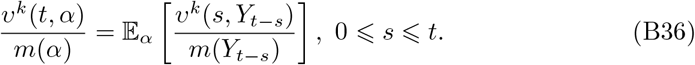

This equality means that the probability of sampling an individual of type *k* among individual with *α* at time *t* is equal to the probability of drawing an individual of type *k* among individuals with trait *y* in the past, at the time *t* − *s* (right hand side). In particular, the previous long time behavior property (B35), shows that the ancestral process converges as *s* → ∞ towards a random variable *Y*_∞_, which admits the density

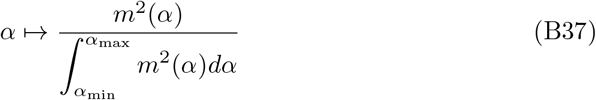

In particular, we show that the most likely common ancestor is a parasitic symbiont with trait *α* = *α*_min_. In addition, on average the trait of the common ancestor is smaller than the mean trait of the actual community (see red and orange curves in Fig. B3). In addition, for small *β*, the mean trait of the common ancestor is mutualistic while for large *β*, it is parasitic (see orange curve in Fig. B3). As a result, we show that the equilibrium is most likely produced by parasitic individuals that mutate and generate the equilibrium distribution. So, mutualism can emerge from parasitism and persist. This behavior persists even for facultative mutualistic plants (Fig. B3c).

## Appendix C Spreading with and without evolution

### C.1 Spreading without evolution

First we investigate the spreading properties of the solution of our problem without variation in trait *α*. So we assume that AMF community is monomorphic: there is only one trait ***α*** in the AMF population. We assume that ***α*** is a real positive constant value between (*α*_min_ and (*α*_max_ + *α*_min_)*/*2, so that the following system admits a constant positive steady state. We investigate the spreading speed of solutions of the following system of equations

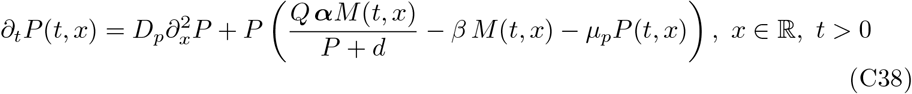

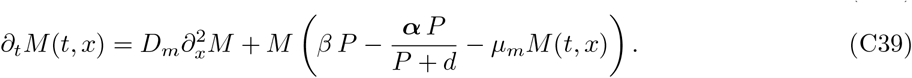

Under, the assumption of Proposition 1, there exists a positive steady state of the model. We aim to investigate the traveling wave solutions of this model, that are solutions of the form *P* (*x* − *ct*) and *M* (*x* − *ct*) where *c* is the spreading speed of the solution and *P* and *M* are profiles that connect the positive stationary state (*P*^∗^, *M*^∗^) to the trivial state (0, 0). Let us remind that the positive steady state satisfies

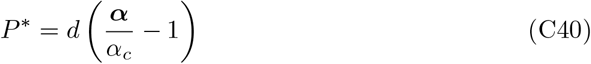

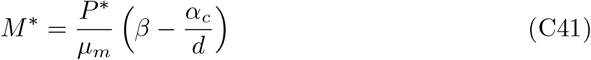

where *α*_*c*_(*β*) is the mutualistic/parasitic threshold defined in (3) and it depends on the parameter *β*.

**Fig. B3.**
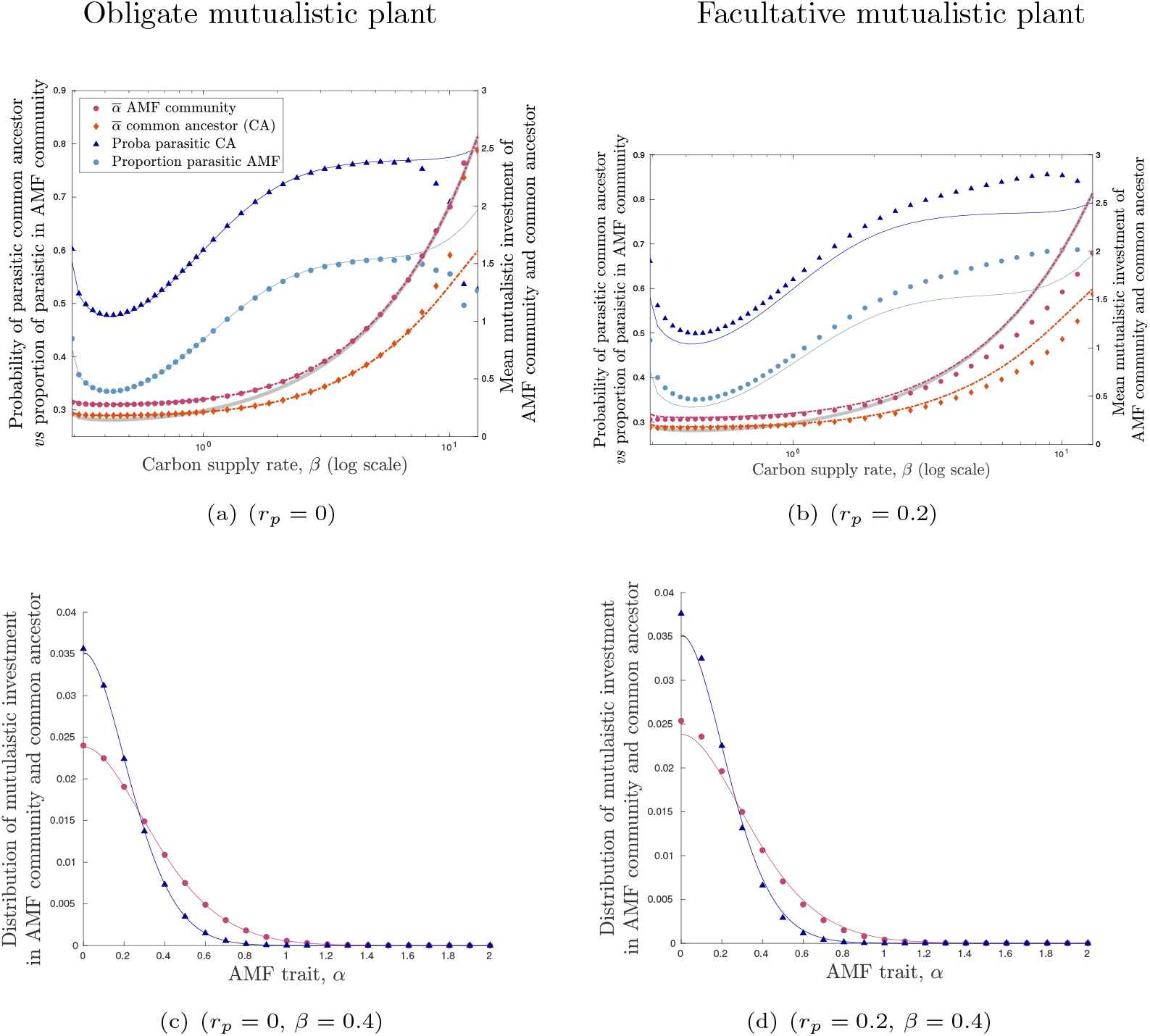
Panel (a-b), Probability of a parasitic common ancestor (dark blue curve and triangles) and mean mutualistic investment of the common ancestor (orange curve and diamonds) compared respectively with the proportion of parasitic symbiont in the AMF community at equilibrium without spatial spread (light blue curve and dots) and the mean mutualistic investment of the community (red curve and dots) associated with obligate mutualistic plant (panel (a)) and facultative mutualistic plant (panel b). Distribution of the mutualistic investment for the common ancestor (dark blue curve and triangles) and the AMF community (red curve and dots) with obligate mutualistic plant (panel (c)) and facultative mutualistic plant (panel d). The curves correspond to the approximations, while the markers correspond to the outcome of the model. For the simulations, the parameters are: *d*_*m*_ = 0.01, *Q* = 6, *μ*_*p*_ = *μ*_*m*_ = 0.3, *β* = 0.72 and *α*_min_ = 0 and *α*_max_ = 5.

To investigate the spreading speed problem, we look at the following equations

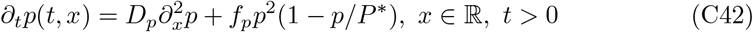

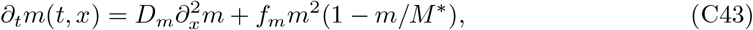

where *f*_*p*_ and *f*_*m*_ are positive constants. We know (Turchin, 1998) that these equations, which are linked only through the positive steady state (*P*^∗^, *M*^∗^), admit a unique traveling wave solutions, that move at a speed *c*_*p*_ and *c*_*m*_ respectively. The speed of spread are given by the following formula

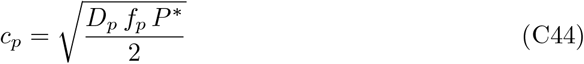

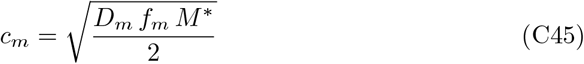

For appropriate constant *f*_*p*_ and *f*_*m*_, the associated solutions *p* and *m* should be sub-solutions of the initial problem (C38). More precisely, we choose *f*_*p*_ and *f*_*m*_ as follows. Let *γ* be a positive constant that we will determine later, and define

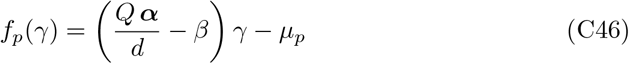

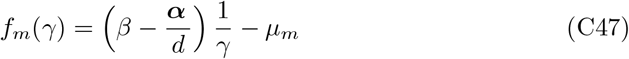

We choose *γ* such that *c*_*p*_(*γ*) = *c*_*m*_(*γ*), so that the solutions *p* and *m* will spread at the same speed

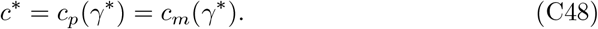

The parameter *γ*^∗^ should be the positive root of the following polynomial function

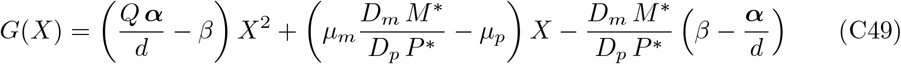

The solution *m* is a super-solution of the initial system if *β* and *γ* are large enough. First, if the solution (*P, M*) of (C38) start from the initial condition (*P* ^0^, *M* ^0^) which is bounded by

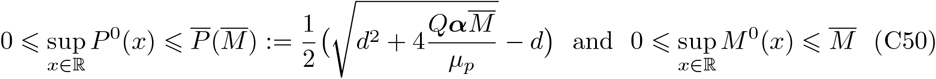

with the constant 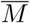such that

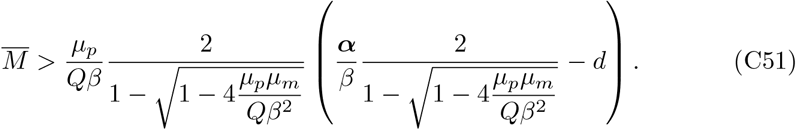

then, the solution (*P, M*) is also bounded for any time *t >* 0.

If *M* is uniformly bounded by 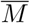 for any positive time *t >* 0 then *P* is also bounded for any time *t >* 0. In this case, a natural super-solution of *P* is a solution 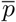 of the following equation

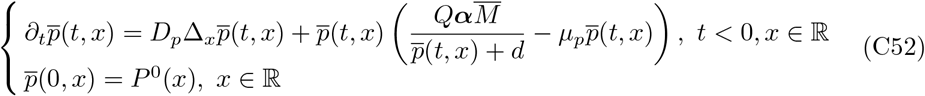

Since, this equation admits the following constant positive steady state

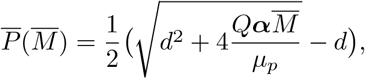

we deduce that this constant is a super-solution of problem (C38) starting from *P* ^0^ and thus from comparison theorem, we deduce that

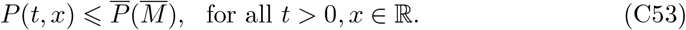

Now, assume by contradiction that *M* is not bounded by 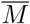, then there exists a positive time *t*_0_ such that 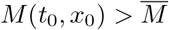 for some location *x*_0_ ∈ ℝ. If we define *T* as follows

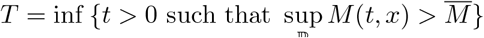

By definition of 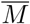 we know that *T >* 0 and we have *T* ⩽ *t* _0_. We can thus apply the same argument as above to show that

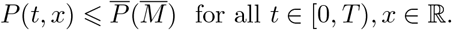

Then, the solution 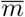 of the following problem:

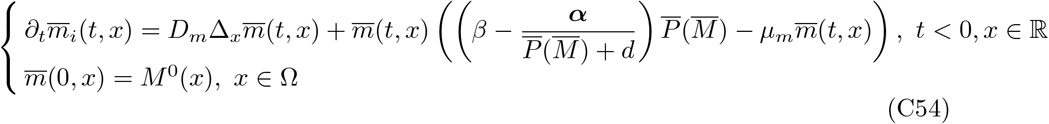

is a super-solution of (C38). In particular, we can show that

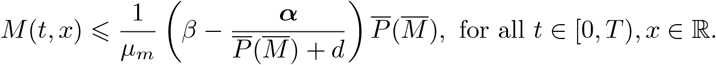

Thus for 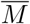 large enough, that is 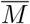 satisfies (C51), we deduce that

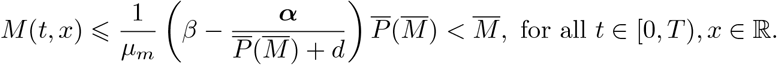

which contradicts the definition of *T* . Thus the solution *M* should be uniformly bounded by 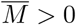 large enough.

Now, let us prove that the ratio *γ*(*t, x*) = *M/P* is uniformly bounded if *D*_*m*_ = *D*_*p*_ = *D*. First, the function *γ* is solution to the following problem

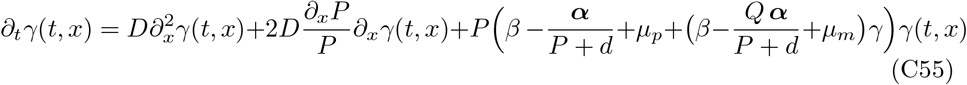

Using the estimates on *P* and *M*, we can show the following estimates for *γ* if *β* is large enough,

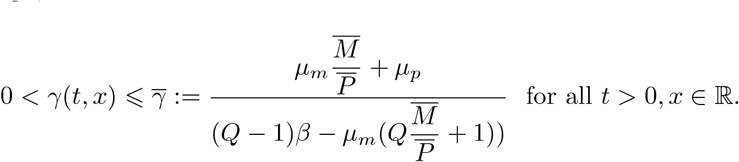

Now let us prove that *m* is a super-solution of the initial problem, starting with appropriate initial condition. Using the estimates on *P, M* and *M/P*, we can show that if *β* large enough, for all *t >* 0, *x* ∈ ℝ and 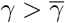, we have

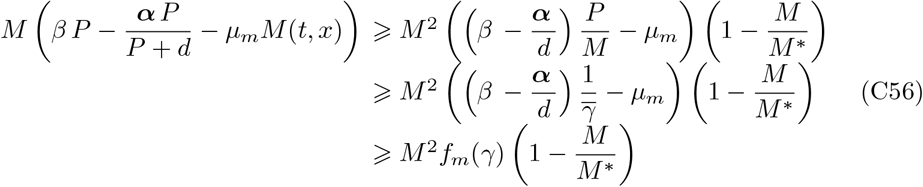

Since *m* is a super-solution of the initial system, the solution (*P, M*) should travel at a speed *c < c*^∗^ for large *β* (Fig. C4).

### C.2 Spreading with evolution

#### Speed of spread

We now look at the spreading sped of the solution with evolution. In this case, we know that the biomass satisfy the following problem

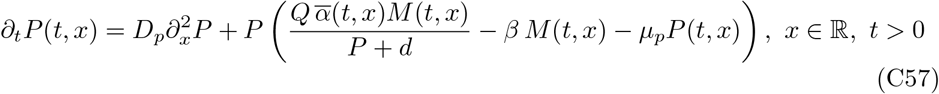

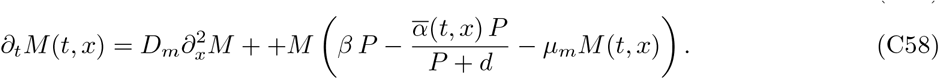

where 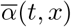 is increasing and satisfies the following property 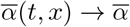 as *t* → + ∞ locally uniformly in space, where 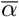 solves the steady state problem (B11)-(B12). From numerical simulation we see that 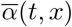 is increasing with respect to the space variable *x*. Under these conditions, we use the previous approximation (C48) to derive the following numerical estimate

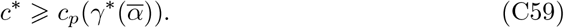

**Fig. C4.**
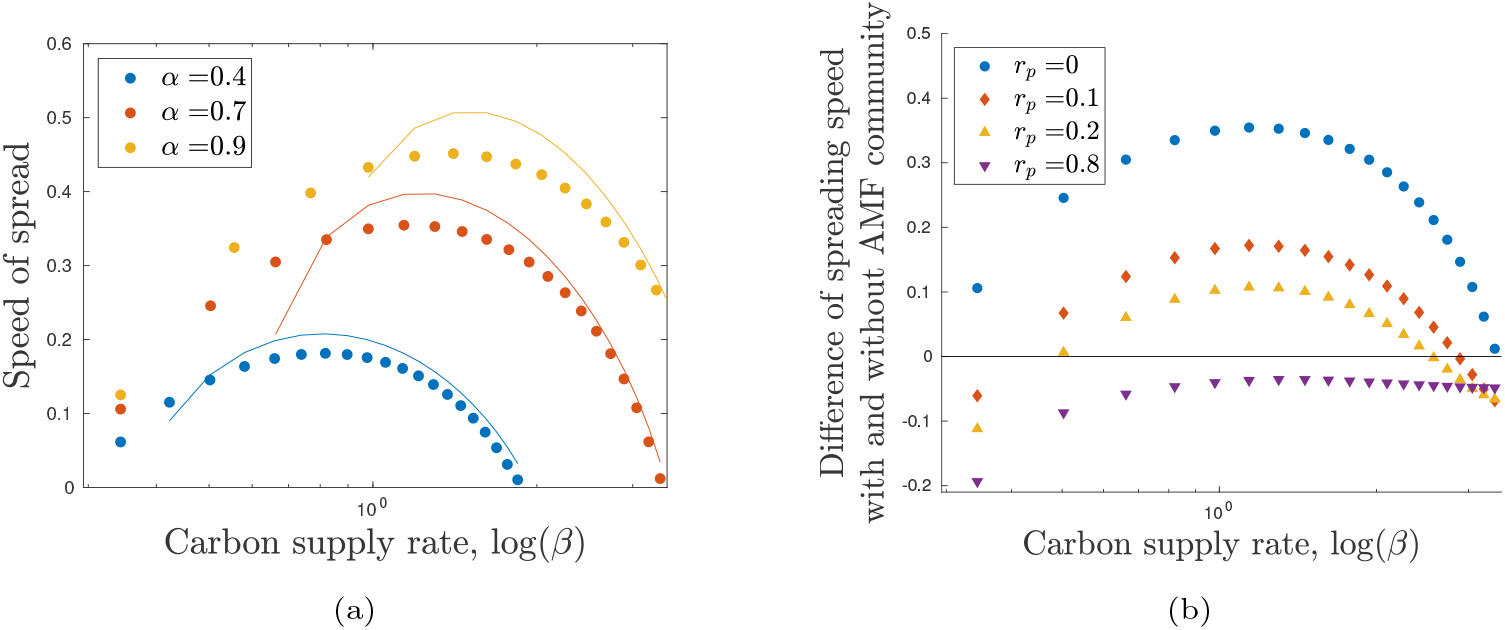
Panel (a), Effect of the carbon supply rate *β* on the speed of spread a plant associated with one AMF with various fixed trait *α* (panel (a)): blue curve (*α* = 0.4), red curve (*α* = 0.7) and orange curve (*α* = 0.9); Panel (b), Difference between the speed of spread of a plant with and without the monomorphic AMF community for various plants with different carbon supply rate and different degree of dependence on the symbiont: obligate mutualistic plants (blue, *r*_*p*_ = 0), facultative mutualistic plants with various intrinsic growth rates, *r*_*p*_ = 0.1 (red), *r*_*p*_ = 0.2 (orange) and *r*_*p*_ = 0.8 (purple). The markers correspond to the speed of the model (1) without evolution *d*_*m*_ = 0) and the curves correspond to the approximation speed (C48). For the simulations, the parameters are: *D*_*p*_ = *D*_*m*_ = 0.1, *Q* = 6, *μ*_*p*_ = *μ*_*m*_ = 0.3.

In this case, the mean mutualistic investment 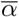 varies over space, and it is more complicated to provide super or sub-solutions for the problem.

#### Fixation probability along the range expansion

Here, we aim to understand how the propagation occurs in space. We first assume that the population of host and symbionts are represented by traveling waves of profile **P**(*x*) and **m**(*x, α*) respectively, which satisfy the following problem

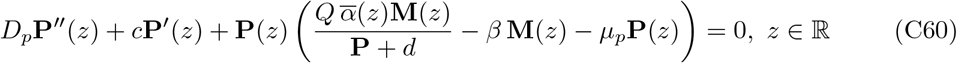

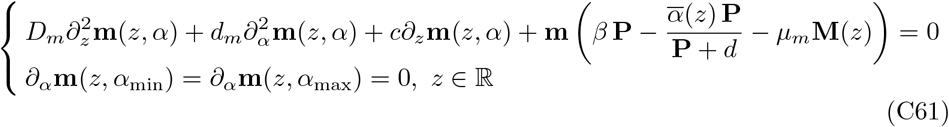

where

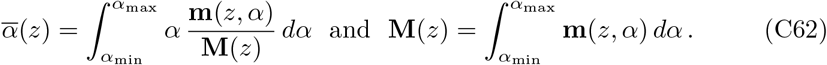

The traveling wave profile satisfy the following boundary conditions

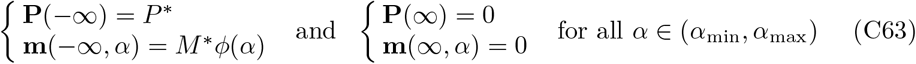

where (*P*^∗^, *M*^∗^, *ϕ*) solves the stationary problem (B11) and (B12)

We use the inside dynamics approach presented in the previous section B.5 and introduced by (Roques et al, 2012). We here label the individuals – hosts or symbionts – according to their initial position in the front, and they transmit their label to their offspring. Since they only differ by their label and their trait for the symbionts, each label *k* ∈ ℕ corresponds to a neutral fraction of density 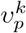 inside the host population **P**(*x* − *ct*) and 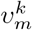 inside the symbiont population **M**(*x* − *ct, α*). Initially, the fractions satisfy

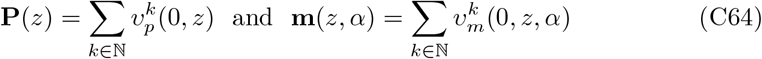

Their dynamics in the moving frame of speed *c*, is described by

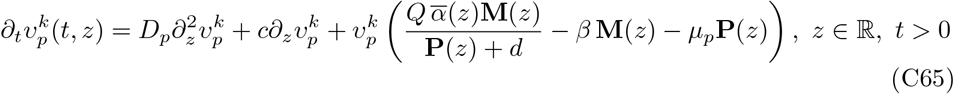

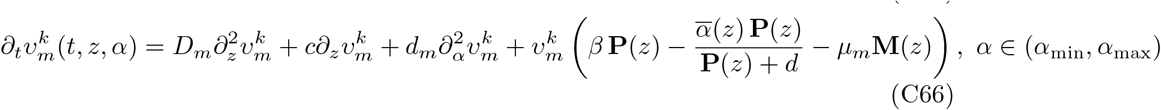

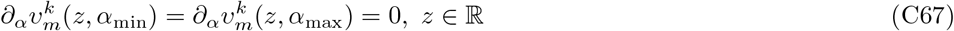

Since we are interested in the effect of initial space location, we can integrate the equation in 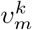 and we obtain that the quantity 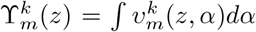 satisfies the following problem

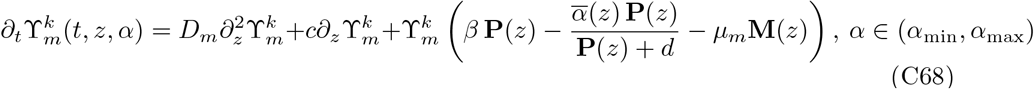

Formally, the solutions 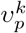 and 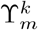 converge uniformly in trait space *α* and locally uniformly in space *z*, toward 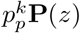 and 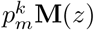 respectively, where the positive constant 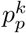 and 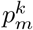 satisfies

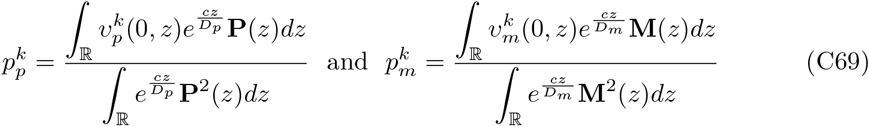

As a result, the fixation probability of host and symbiont individuals starting at location *x*, that is the probability that their descendants become prevalent at the leading edge of the travelling wave, is given by, respectively, *PF*_*p*_(*x*) and *PF*_*m*_(*x*), which are defined by

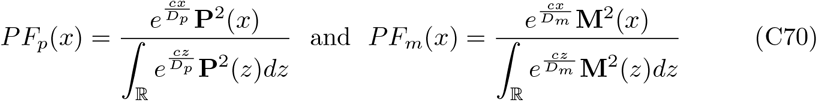

Since the traveling profiles **P** and **M** decay at different rates (Fig. 3), the fixation probability attains is maximum at different location. In particular, we see that the plant at the leading edge have a higher probability of fixation than the symbiont at the same location. Thus plant at the leading edge mainly come from this edge, while the symbiont come more from the core of the population (Fig. 3).

